# Genome-wide DNA methylation patterns harbor signatures of hatchling sex and past incubation temperature in a species with environmental sex determination

**DOI:** 10.1101/2022.05.03.490459

**Authors:** Samantha L. Bock, Christopher R. Smaga, Jessica A. McCoy, Benjamin B. Parrott

**Author notes:** Corresponding author: Samantha Bock.

## Abstract

Conservation of thermally sensitive species depends on monitoring organismal and population-level responses to environmental change in real time. Epigenetic processes are increasingly recognized as key integrators of environmental conditions into developmentally plastic responses, and attendant epigenomic datasets hold potential for revealing cryptic phenotypes relevant to conservation efforts. Here, we demonstrate the utility of genome-wide DNA methylation (DNAm) patterns in the face of climate change for a group of especially vulnerable species, those with temperature-dependent sex determination (TSD). Due to their reliance on thermal cues during development to determine sexual fate, contemporary shifts in temperature are predicted to skew offspring sex ratios and ultimately destabilize sensitive populations. Using reduced-representation bisulfite sequencing, we profiled the DNA methylome in blood cells of hatchling American alligator (*Alligator mississippiensis*), a TSD species lacking reliable markers of sexual dimorphism in early life-stages. We identified 120 sex-associated differentially methylated cytosines (DMCs; FDR < 0.1) in hatchlings incubated under a range of temperatures, as well as 707 unique temperature-associated DMCs. We further developed DNAm-based models capable of predicting hatchling sex with 100% accuracy and past incubation temperature with a mean absolute error of 1.2°C based on the methylation status of 20 and 24 loci, respectively. Though largely independent of epigenomic patterning occurring in the embryonic gonad during TSD, DNAm patterns in blood cells may serve as non-lethal markers of hatchling sex and past incubation conditions in conservation applications. These findings also raise intriguing questions regarding tissue-specific epigenomic patterning in the context of developmental plasticity.

## Introduction

Climate change threatens species persistence in wild populations via its influence on key demographic parameters. In ectotherms, shifting temperature regimes impact juvenile recruitment (Laloë et al., 2017; Noble et al., 2018; Pruett & Warner, 2021; Schwanz et al., 2010), adult fecundity (Crews et al., 1998; Wang et al., 2016; Warner & Shine, 2008), and, in some cases, population sex ratios (Bock et al., 2020; Janzen, 1994; Jensen et al., 2018; Reneker & Kamel, 2016). Fortunately, efforts to predict organismal and population level responses to changing environmental conditions are rapidly advancing with improvements in the precision and geographic resolution of global climate models and their integration with empirical data from physiological, ecological, and evolutionary studies (Carlo et al., 2018; Laloë et al., 2014; Mitchell et al., 2008; Riddell et al., 2018; Seebacher et al., 2015). Yet, one major challenge moving forward will be ground truthing these predictions to adapt conservation strategies accordingly. Genomic and epigenomic datasets represent powerful resources for conservationists in these efforts (Formenti et al., 2022; Layton & Bradbury, 2021; Waldvogel et al., 2020). The use of genomic tools in conservation is already widespread, allowing for assessments of population structure and connectivity (Wright et al., 2020), inbreeding depression (Dussex et al., 2021; Kardos et al., 2016), contemporary evolutionary responses to environmental change (Bi et al., 2019; Catullo et al., 2019), and adaptive capacities to cope with future change (Bay et al., 2018; Flanagan et al., 2018; Harrisson et al., 2014; Layton et al., 2021). The potential of epigenomic tools, on the other hand, is only just beginning to be realized. DNA methylation (DNAm), the covalent addition of a methyl group to a cytosine base, is one of the best-studied epigenetic marks, and variation in DNAm within and across individuals harbors key biological and ecological insights (Angers et al., 2010; Schübeler, 2015; Ziller et al., 2013). In the face of a rapidly shifting global climate, variation in genomic DNAm patterns has the potential to reveal cryptic organismal traits relevant to conservation and may provide insight into the environmental conditions from which they arose.

The DNA methylome exists at the interface of the genome and the environment, operating in a coordinated manner with other epigenetic modifications to influence gene regulation (Feil & Fraga, 2012; Luo et al., 2018; Zhu et al., 2016). Genome-wide patterns of DNAm can be stably maintained through cell division (Bird, 2002), and thus reflect interactions between the underlying genetic sequence and both past and present environmental conditions. These features have led biomedical fields to turn to DNAm-based biomarkers to gain insight into biological aging (Horvath, 2013; Horvath & Raj, 2018), contaminant exposure (Cardenas et al., 2017; Green et al., 2016; Gruzieva et al., 2017; Joubert et al., 2016), and the developmental origins of adult health and disease (Felix & Cecil, 2019; Heijmans et al., 2008). More recently, the ecological sciences have followed suit, harnessing the potential of the DNA methylome to understand life-history variation (Anderson et al., 2021; Cayuela et al., 2021; de Paoli-Iseppi et al., 2017; Lindner et al., 2021; Parrott & Bertucci, 2019), organismal health (Crossman et al., 2021; Hu et al., 2018; Lea et al., 2016), exposure to environmental stressors (Bertucci et al., 2021; Guillette et al., 2016; Liew et al., 2018; Mäkinen et al., 2021; Parrott, Bowden, et al., 2014), and developmental plasticity (Laubach et al., 2019; Navarro-Martín et al., 2011; Parrott, Kohno, et al., 2014; vonHoldt et al., 2021). Collectively, these studies indicate that DNAm patterns at discrete loci in non-lethally sampled tissues (e.g., blood) provide insight into fundamental aspects of an individual’s biology, health status, and past environmental experience that would otherwise be nearly impossible to measure.

These observations are particularly relevant for thermally sensitive species facing climate-induced population collapse. Among these vulnerable taxa are species with temperature-dependent sex determination (TSD). Due to their reliance on temperature cues during incubation to determine offspring sex, species with TSD, including many turtles, all crocodilians, and some fish, are predicted to experience highly skewed population sex ratios as a result of climate change (Bock et al., 2020; Janzen, 1994; Laloë et al., 2014; Mitchell et al., 2008, 2010; Ospina-Alvarez & Piferrer, 2008; Reneker & Kamel, 2016). Such skews in sex ratios are already being detected in wild populations of TSD species (Jensen et al., 2018) and may exert lasting adverse impacts on population dynamics given the long generation times of many TSD species (Böhm et al., 2016). Despite these observations, efforts to actively monitor cohort sex ratios in the field remain limited. This is in part due to the lack of reliable, non-lethal methods of sexing individuals at early life stages. Since hatchlings do not tend to exhibit sexually dimorphic morphology, specialized training is required to determine hatchling sex (Allsteadt & Lang, 1995), and genetic markers of sex (i.e., sex-linked genes or alleles) are non-existent in TSD species (or may not be indicative of phenotypic sex in the case of temperature-induced sex-reversal (Holleley et al., 2015)). DNAm patterns in blood represent a novel avenue by which those tasked with managing populations of TSD species under climate change might gain insight into ongoing sex ratio shifts as well as changes in field incubation conditions.

The DNA methylome appears to integrate temperature cues into sexually dimorphic phenotypic trajectories during TSD, though the details of this process have yet to be resolved at a genome-wide scale. During development, gonadal tissues acquire sex-associated DNAm patterns at the promoters of genes with conserved roles in vertebrate sex determination in response to incubation temperature (Matsumoto et al., 2016; Navarro-Martín et al., 2011; Parrott, Kohno, et al., 2014). It remains unknown whether somatic tissues that lack direct roles in sex determination also acquire similar sex- or temperature-associated patterns. However, a few lines of evidence from species with genotypic sex determination support this possibility. Several studies have identified autosomal sex-associated variation in DNAm patterns across species differing in sex-determination system in somatic tissues (Gatev et al., 2021), including blood (Caizergues et al., 2021; Janowitz Koch et al., 2016), muscle (Wan et al., 2016), and liver (Hu et al., 2019; Zhuang et al., 2020). Some sex-biased autosomal DNAm patterns have even been shown to be established during development and stable throughout the life course (Gatev et al., 2021).

Though such differences in DNAm patterns tend to be more subtle in autosomal regions of the genome compared to sex chromosomes, these findings suggest the DNA methylomes of somatic tissues may harbor sex-associated patterns in TSD species. Further, environmental influences on DNAm patterns established during development can persist in the post-natal period even after the environmental cue has passed (Gavery et al., 2019; vonHoldt et al., 2021). Developmental thermal cues can exert lasting influences on the DNA methylome (Anastasiadi et al., 2021; Metzger & Schulte, 2017), raising the possibility that DNAm patterns in hatchlings might provide information about past incubation conditions.

In this study, we investigate sex- and temperature-associated variation in the DNA methylomes of hatchling blood cells in the American alligator, a species with TSD. DNAm dynamics in blood cells are examined in the context of other sex-linked biological correlates such as circulating levels of sex steroids and gonadal DNAm patterns established during TSD. We further use these DNAm patterns to develop models that predict hatchling sex and past incubation temperature based on the methylation status of a subset of loci. Using these predictive models, we assess the potential utility of DNAm-based markers in the conservation of species with TSD under rapid environmental change. Due to the unique temperature-by-sex ratio reaction norm of the alligator, this species serves as a particularly insightful model to identify sex- and temperature-associated DNAm patterns. As in all other crocodilians, females are produced at both low and high incubation temperatures, while males are produced at intermediate incubation temperatures (Lang & Andrews, 1994). From the perspective of experimental design, this pattern of TSD allows us to disentangle DNAm variation linked to sex from that linked to temperature through the inclusion of females from both the low and high ranges of the reaction norm. The shape of the crocodilian temperature-by-sex ratio reaction norm also yields greater uncertainty in predictions of sex ratio shifts in response to climate change (Bock et al., 2020). Thus, monitoring sex ratios in the field is especially pertinent for these species and our results hold unique relevance for future conservation.

## Materials and Methods

### Incubation experiment and sample collection

In June 2013, seven clutches of *A. mississipiensis* eggs were collected from Lake Apopka, FL and transported to Hollings Marine Laboratory in Charleston, SC (McCoy et al., 2016). Detailed methods regarding the incubation experiment can be found in the associated report, which sought to decipher temperature-dependent inter- and intra-sexual variation in gonadal gene expression (McCoy et al., 2016). Briefly, eggs were kept in damp sphagnum moss within commercial incubators (Thermo Scientific, Forma Environmental Chambers, model #3920). One embryo from each clutch was staged according to Ferguson developmental staging criteria (Ferguson, 1985). Viable eggs were maintained at 30°C until stage 19, the opening of the canonical thermosensitive period (Lang & Andrews, 1994), after which eggs from each clutch were distributed across four incubation temperatures (30°C, 32°C, 33.5°C, 34.5°C), where they were maintained until hatch. All animal protocols were approved by the Medical University of South Carolina’s Institutional Animal Care and Use Committee (AUP # 3036).

Following hatch, alligator neonates were measured to assess morphometric traits including mass, snout-vent length, and total length. At seven days post-hatch, body morphometric measurements were repeated, and a blood sample was taken from the postcranial sinus. Animals were then euthanized with a lethal dose of pentabarbital (Sigma Aldrich) and necropsied to collect the whole brain and gonad-adrenal-mesonephros (GAM) complex. Fresh tissues were preserved in RNAlater (Life Technologies) and kept at -80°C. Whole blood samples were stored in lithium heparin Vacutainer tubes (BD) on ice prior to centrifugation at 1500 rcf for 20 min. Plasma was isolated and RNAlater (Life Technologies) was added to the residual pelleted blood cells. Plasma was stored at -80°C and blood cells in RNAlater were stored at - 20°C. Plasma concentrations of 17ß-estradiol (E2) and testosterone (T) were quantified via radioimmunoassays as described in (McCoy et al., 2016).

### Temperature-by-sex reaction norm, sex ratio assessment, and sample selection

To place the incubation treatments and resulting hatchling sex ratios of the present study in the broader context of historical incubation experiments, we gathered data from 14 studies in the American alligator employing constant incubation temperatures and quantifying hatchling sex ratios (Table S1). An equation for the temperature-by-sex ratio reaction norm in this species was then fit with this data using logistic regression with a quadratic term for temperature (Figure 1).

**Figure 1.**
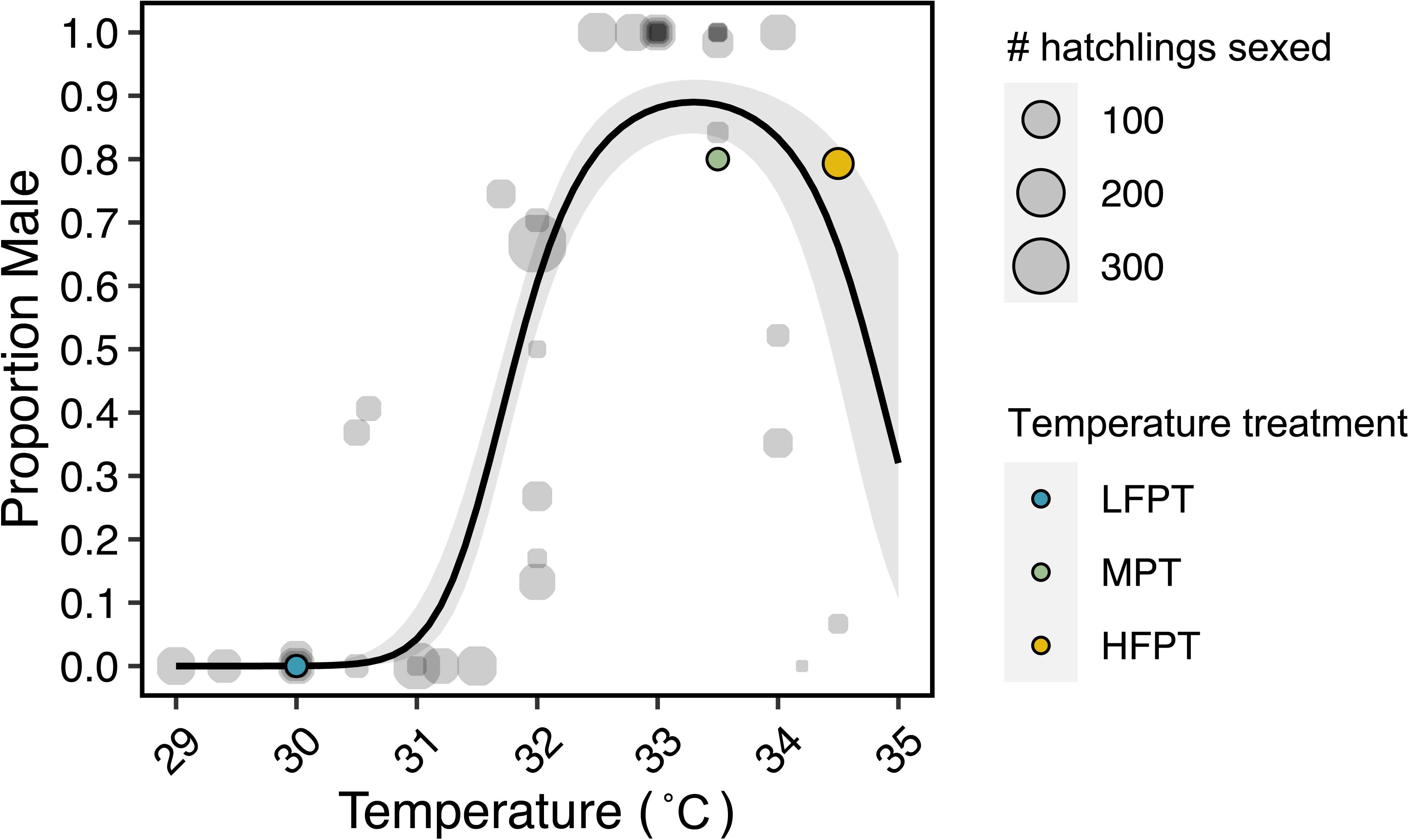
Incubation temperature treatments in the context of the American alligator temperature-by-sex ratio reaction norm. Each point corresponds to the resulting sex ratio from a single incubation treatment from one of fourteen experiments. Points corresponding to treatments and resulting sex ratios used in this study are colored according to their temperature (30°C = LFPT, blue; 33.5°C = MPT, green; 34.5°C = HFPT, yellow). Point size is proportional to the number of hatchlings that were sexed. Solid line indicates fit of logistic model of sex ratio with quadratic term for temperature, and shaded area indicated 95% confidence interval.

In the present study, hatchling sex was assigned based on gonadal expression of *AMH* and *CYP19A1* in (McCoy et al., 2016). These transcripts exhibit robust sex-biased expression patterns in hatchling gonads, and in a previous study, sex assignment based on expression of these genes was confirmed by histology in 98.7% (213/216) of samples (Kohno et al., 2015). Consistent with previous reports of the alligator temperature-by-sex reaction norm (Lang & Andrews, 1994), eggs incubated at 30°C yielded 100% female hatchlings (n = 19), while eggs incubated at 33.5°C and 34.5°C yielded primarily male hatchlings (80% male, 16/20; and 79.3% male, 46/58 respectively; Figure 1). In order to identify sexually dimorphic DNAm patterns in blood cells, samples from eight females incubated at 30°C, eight males incubated at 33.5°C, four females incubated at 34.5°C, and four males incubated at 34.5°C were selected for reduced representation bisulfite sequencing (RRBS) (Table 1). Clutch identity was also accounted for in sample selection, ensuring that clutch representation (N = 5 clutches included) was balanced across incubation temperatures and sexes.

**Table 1.** Study design and sampling scheme.

| Temperature (°C) | Clutch |  |  |  |  | Total |
| --- | --- | --- | --- | --- | --- | --- |
|  | AP-03 | AP-04 | AP-05 | AP-06 | AP-08 |  |
| 30 | 2 | 2 | 1 | 1 | 2 | 8 |
| 33.5 | 2 | 2 | 1 | 1 | 2 | 8 |
| 34.5 (F) | 2 | 1 | 0 | 1 | 0 | 4 |
| 34.5 (M) | 2 | 1 | 0 | 1 | 0 | 4 |

| Temperature (°C) | Clutch |  |  |  |  |  |
| --- | --- | --- | --- | --- | --- | --- |
|  | AP-03 | AP-04 | AP-05 | AP-06 | AP-08 | Total |
| 30 | 2 | 2 | 1 | 1 | 2 | 8 |
| 33.5 | 2 | 2 | 1 | 1 | 2 | 8 |
| 34.5 (F) | 2 | 1 | 0 | 1 | 0 | 4 |
| 34.5 (M) | 2 | 1 | 0 | 1 | 0 | 4 |

### Nucleic acid isolation and quantification

DNA was isolated from blood cells using a modified column approach based on the SV total RNA isolation system (Promega). Frozen samples were thawed on ice. To remove RNAlater from the preserved cells, 500 μl of phosphate-buffered saline (PBS) was added to a 200 μl aliquot of each sample and samples were centrifuged at 5000 rcf for 15 minutes at 4°C (and for an additional 5 min if a cell pellet did not form). Pelleted cells were washed once more with 500 μl of PBS and nucleic acids were isolated according to the protocol reported in (Bae et al., 2021; Appendix S1). Resulting DNA was eluted in 60 µl TE buffer (pH 8.0). DNA concentrations and qualities were quantified using a Qubit fluorometer (Thermo Fisher) and NanoDrop spectrometer (Thermo Fisher) respectively. Extracted DNA was stored at -80°C until library preparation.

### Reduced-representation bisulfite sequencing

Reduced-representation bisulfite sequencing (RRBS) libraries were constructed with 200 ng input DNA from each sample (n = 24; Table 1) using the Premium RRBS Kit (Diagenode, Cat. No. C02030032) according to manufacturer instructions. Unmethylated and methylated spike-in control sequences are included within the commercial kit to allow for assessment of bisulfite conversion efficiency, which in this experiment exceeded 99% for all samples (mean ± SD = 99.57± 0.25). Size selection and clean-up steps were performed with AMPure XP Beads (Beckman Coulter), and bisulfite-converted libraries were subjected to between 13-15 amplification cycles in the final PCR. Resulting libraries were stored at -80°C prior to submission to the Georgia Genomics and Bioinformatics Core (GGBC; University of Georgia, Athens GA) for sequencing. Quality of the libraries was assessed using an Agilent Fragment Analyzer, and pooled libraries were sequenced on a single Illumina NextSeq 500 High Output flowcell (75bp, single-end) with 20% Illumina PhiX control. Sequencing generated a total of 300 million reads, with 94% meeting or exceeding the Phred score quality threshold of >30. Raw reads were assessed for quality using FastQC (version 0.11.8) (https://www.bioinformatics.babraham.ac.uk/projects/fastqc/) and MultiQC (version 1.8) (Ewels et al., 2016).

### Quality trimming, read alignment, and methylation data processing

Raw reads were trimmed to remove adapter contamination and low-quality bases (Phred score < 20) using TrimGalore! (version 0.6.5) with the ‘--rrbs’ option and a minimum overlap with the adapter sequence of 1 bp (‘-stringency 1’) (https://www.bioinformatics.babraham.ac.uk/projects/trim_galore/). After quality trimming, 207 million reads remained across all samples, with a mean (± SD) of 8.62 ± 1.13 million reads per sample. Alignment of trimmed reads to the latest American alligator genome assembly (ASM28112v4) (Rice et al., 2017) and methylation calling was performed with Bismark (version 0.22.3) (Krueger & Andrews, 2011) and Bowtie 2 (version 2.4.1) (Langmead & Salzberg, 2012) using default parameters. Alignment rates ranged from 61.8% to 66.0% (mean 63.6%; SD 1.25%). Resulting BAM files were sorted and indexed with SAMtools (version 1.10) (Li et al., 2009), then read into R (version 4.0.0) (R Core Team, 2021) with the “processBismarkAln” function in the methylKit package (version 1.14.2; Bioconductor version 2.48.0) (Akalin et al., 2012). Analyses were limited to cytosines in a CpG context, as most cytosine methylation in vertebrate genomes occurs in this context (Feng et al., 2010). Using the methylKit and GenomicRanges packages (version 1.40.0) (Lawrence et al., 2013), reads on both strands of a CpG site were merged and loci were filtered to remove those in the 99.9th percentile of coverage to account for PCR bias. Loci covered by less than 5 reads in 50% of samples per treatment group (temperature-sex combination) were also removed. Read coverage was then normalized across samples within methylKit. Prior to differential methylation analysis, constitutively hyper- and hypo-methylated CpG sites (>95% or <5% methylation across all samples) were removed, as well as invariable CpG sites defined by the “nearZeroVar” function within the caret package (version 6.0-86) (Kuhn, 2008).

### Differential methylation analysis

Differentially methylated cytosines (DMCs) were identified via five main comparisons. To identify ‘sex-associated’ DMCs, all females were compared to all males (FvM) and 34.5°C females were compared to 34.5°C males (F^34.5^vM^34.5^). To identify ‘temperature-associated’ DMCs, pairwise comparisons of hatchlings from each temperature treatment were conducted (30v33.5, 30v34.5, 33.5v34.5). If a cytosine was identified as differentially methylated in both the sex- and temperature-based differential methylation analyses, the site was considered ‘sex-associated’. Differential methylation was modeled in methylKit using logistic regression and overdispersion of variance was corrected for according to (McCullagh & Nelder, 1989). A cytosine was defined as differentially methylated with respect to the focal comparison if it passed a 10% Benjamini-Hochberg False Discovery Rate (FDR) threshold. Resulting ‘sex-associated’ DMCs were categorized based on whether percent methylation was greater in females or males, while ‘temperature-associated’ DMCs were categorized based on whether percent methylation increased or decreased with incubation temperature. We also assessed temperature-associated DNAm patterns by performing Spearman correlations between incubation temperature and percent methylation for all covered cytosines using the “corAndPvalue” function within the WGCNA package (version 1.70-3) (Langfelder & Horvath, 2008). To correct for multiple comparisons, an FDR approach was applied using the “p.adjust” function in R. For this correlation analysis, as well as further correlation analyses and predictive analyses described later, percent methylation values were imputed for samples with missing values (i.e., locus without sufficient read coverage in the sample) using a k-nearest neighbor approach implemented with the “impute.knn” function in the impute package (version 1.62.0) under default parameters (k = 10) (Troyanskaya et al., 2001). The absolute Spearman correlation coefficients with temperature for loci identified as temperature-associated DMCs were compared to those of loci not identified as temperature-associated DMCs using a non-parametric Mann-Whitney U test.

### Annotation of differentially methylated loci

Differentially methylated cytosines were annotated according to their gene context (i.e., promoter, exon, intron, or intergenic) using the alligator genome annotation (GCF_000281125.3, annotation release 102) within the GenomicRanges package in R. If a cytosine overlapped with multiple gene contexts, the site was defined according to the following precedent: promoter > exon > intron > intergenic. The alligator gene annotation table from NCBI was used within GenomicRanges to determine the closest gene to each cytosine in the dataset using the “distancetoNearest” function. Gene ontology (GO) term enrichment analysis was performed for unordered gene lists associated with DMCs identified as temperature-associated, sex-associated across all individuals, and sex-associated across all individuals or within 34.5°C individuals using g:Profiler (version e105_eg52_p16_e84549f) (Raudvere et al., 2019). Separate GO analyses were performed for gene lists corresponding to DMCs overlapping promoters and gene lists corresponding to DMCs overlapping promoters or gene bodies. GO terms were considered significantly enriched if they passed a Benjamini-Hochberg FDR threshold of 10% against a custom background composed of genes associated with all analyzed cytosines. Any genes lacking annotations (i.e., uncharacterized ‘LOC’ genes) were excluded from analysis.

The regulatory potential and functional role of individual CpG sites is often linked to local CpG density, especially in the context of CpG ‘islands’ in mammals (Weber et al., 2007). Thus, to better understand the relevance of DMCs identified in our analyses, we annotated these loci with respect to CpG density via two complementary approaches. First, we predicted the locations of CpG islands (CGIs) in the alligator genome using the program ‘cpgplot’ (EMBOSS version 6.6.0; parameters: -window 100, -minlen 200, -minoe 0.6, -minpc 50) (Rice et al., 2000). The locations of putative CGIs were then used to define the locations of CpG shores (± 2000 bp of CGIs) and CpG shelves (± 2000 bp of shores) using BEDtools (version 2.26.0) (Quinlan & Hall, 2010). Genomic regions not included in CpG islands, shores, or shelfs were defined as “open sea” regions. Overlap of DMCs with CGIs, shores, shelves, and open sea regions was determined within GenomicRanges in R. In addition to this discrete approach of defining CGIs in the alligator genome, we also assessed the local CpG density around DMCs on a more continuous scale. To do this, the entire genome was tiled into non-overlapping 1kb windows using BEDtools. The CpG density of each tile was then quantified by determining its overlap with all the CG dinucleotides in the genome. Locations of all the CG dinucleotides in the alligator genome were determined with Python (version 3.8.2) using the “fastaRegexFinder.py” script (https://github.com/dariober/bioinformatics-cafe). DMCs were then defined by the CpG density of their overlapping tile (with CpG density being binned into 20 CpG/kb bins).

Differentially methylated cytosines were also characterized according to the overall percent methylation at these loci. Mean percent methylation across all samples was determined for all cytosines analyzed, then loci were categorized into one of five bins (mean percent methylation = 0-20%, 20-40%, 40-60%, 60-80%, or 80-100%). To determine enrichment or depletion of differentially methylated cytosines within genomic regions of interest, Fisher’s exact tests were conducted to compare DMC annotations with the equivalent annotations of all the cytosines analyzed (i.e., those meeting coverage and filtering requirements; N = 462,236 – (# DMCs)). The cytosines included in our analyses were also compared to all the CG dinucleotides in the alligator genome (N = 21,480,261 – (# covered CpGs)). Fisher’s exact tests followed by FDR correction were conducted for each annotation category: gene context, CGI context, CpG density context, and overall percent methylation context.

### Potential regulation by circulating sex steroids

To investigate the potential contribution of circulating sex steroid hormones towards observed differential methylation patterns, we performed Spearman correlations between percent methylation for all covered cytosines and plasma E2 using the “corAndPvalue” in the WGCNA package. We also tested for Spearman correlations between percent methylation and plasma testosterone. Correlations were tested with the imputed percent methylation dataset, and correction for multiple comparisons was performed using an FDR approach. The absolute Spearman correlation coefficients with E2 and testosterone for loci identified as sex-associated DMCs were compared to those of all other covered loci using non-parametric Mann-Whitney U tests. The locations of putative estrogen-response elements (EREs) and androgen-response elements (AREs) in the alligator genome were predicted using the tool PWMscan, a genome-wide position weight matrix (PWM) scanner (Ambrosini et al., 2018), with the motifs for ESR1 (MA0112.3) and AR (MA0007.3) from the JASPAR Core 2020 vertebrate motif library. Putative EREs and AREs meeting a significance threshold of p < 0.000002 (ERE: n = 7,225; ARE: n = 6,149) were used for further analysis. Distance to the nearest ERE and ARE was determined for each DMC, covered CpG site, and CpG in the alligator genome using GenomicRanges in R. Then the proportion of loci falling within 1 kb, 2.5 kb, and 5 kb of an ERE and ARE was determined for each CpG category. The proportion of DMCs identified as sex-associated across all individuals, DMCs identified as sex-associated across all individuals or within 34.5°C, and DMCs identified as temperature-associated within each distance category were compared to the proportion of analyzed CpG sites within each distance category using Fisher’s exact tests followed by FDR correction. Further, the proportion of analyzed CpG sites within each distance category was compared to the proportion of all the CpG sites in the alligator genome within each distance category.

The initial incubation experiment from which we derived these samples, (McCoy et al., 2016), did not detect any significant relationships between plasma sex steroid concentrations and sex or incubation temperature. To confirm this result in the subset of samples included in the present study, we fit generalized linear mixed effect models for E2 and testosterone concentrations with fixed effects of incubation temperature and sex, and a random effect of clutch using the lme4 package in R (Bates et al., 2015). Log-ratio tests were conducted to test which explanatory variables improved model fit. Variables that did not improve model fit were subsequently excluded from the model (according to a significance threshold of p < 0.05), and p-values for variables included in the final models were derived from the lmerTest package, using Satterthwaite’s degrees of freedom method (Kuznetsova et al., 2017).

### DNA methylation-based predictor of sex and incubation temperature

To develop a DNAm-based model to predict hatchling sex, we divided our samples into a training set of 20 individuals and a test set of four individuals. The test set was composed of one randomly selected individual from each temperature-sex combination. Loci exhibiting differential methylation with respect to sex across all individuals in the training set were identified as described previously, however in this case, cytosines were considered differentially methylated if they passed a more stringent FDR threshold of 5%. We then identified the subset of cytosines that were differentially methylated with respect to sex within the training set and across the entire dataset. The imputed percent methylation matrix for these loci was then modeled using penalized binomial logistic regression with elastic net regularization (alpha = 0.5) in the R package glmnet (version 4.1-3) to identify sites for which methylation status was most predictive of sex in the training set (Zou & Hastie, 2005). In this modeling approach, the coefficients of loci that are weak predictors of sex are reduced to zero, thereby eliminating them from the model. The strength of the penalty on model coefficients is controlled by the tuning parameter lambda which is selected to minimize mean-squared error during internal n-fold cross-validation (n-fold = 5). The performance of the model was then assessed by the correct assignment of sex for individuals in the test set.

To develop a DNAm-based predictive model for incubation temperature, a similar approach was taken to the one described for developing a predictive model of sex. Loci exhibiting differential methylation with respect to temperature were used as input for the predictor selection. In this case, we fit a penalized linear regression with elastic net regularization. Model fit on the training set and performance on the test set was evaluated through the R-squared of the relationship between actual and predicted incubation temperature and the absolute error of the predicted incubation temperatures.

### Comparison of DNA methylation patterns between blood and embryonic gonad

To place the temperature-associated DNAm patterns of hatchling blood cells in the broader context of global epigenetic patterning during TSD, we utilized an additional dataset consisting of RRBS data from alligator embryonic gonads sampled just after the closing of the thermosensitive period in development. With this data, we assessed the extent to which genome-wide DNAm responses to incubation temperature differ between somatic (i.e. blood) and gonadal tissues. Briefly, seven clutches of alligator eggs (N = 315) were collected from wild nests in June 2018 at the Lake Woodruff National Wildlife Refuge (De Leon Springs, FL) under permits granted by the Florida Fish and Wildlife Conservation Commission (permit # SPGS-18-33). Following transport to the Savannah River Ecology Laboratory (Jackson, SC), a representative embryo from each clutch was staged according to Ferguson developmental staging criteria (Ferguson, 1985). Viable eggs were maintained in damp sphagnum moss within commercial incubators (Percival Scientific, model I36NLC) at 32°C, a temperature known to produce mixed sex ratios, until stage 15 upon which eggs from each clutch were distributed across four incubation temperatures (29°C, 32°C, 33.5°C, 34.5°C). Embryos were dissected at stage 26, just after the closing of the thermosensitive period when sexual fate has been irreversibly determined, and GAM complexes were preserved in RNAlater (Life Technologies) and kept at -80°C. Gonads were subsequently dissected from adrenal-mesonephros tissue and DNA was extracted, quantified, and assessed for quality according to the same protocol described previously (modified SV total RNA isolation system; Promega). RRBS libraries were prepared for 23 samples with 200 ng genomic DNA according to the protocol described previously (Diagenode Premium RRBS kit) and kept at -80°C until submission to the GGBC for sequencing. Quality of the libraries was assessed using an Agilent Fragment Analyzer, and pooled libraries were sequenced on a single Illumina NextSeq 500 High Output flowcell (75bp, single-end). Sequencing generated a total of 251 million reads, with 93% meeting or exceeding the Phred score quality threshold of >30. Raw reads were processed through the same bioinformatic pipeline as described previously. One sample was excluded from further analysis due to aberrant genomic coverage. Temperature-associated DMCs were then identified in the gonad through pairwise comparisons of individuals from each temperature group using logistic regression with correction for overdispersion as described previously (FDR < 0.1).

To compare DMCs identified in the blood and gonad, we subset each dataset to only include the cytosines that met coverage and filtering requirements in both tissues (N = 181,834). We compared the proportion of shared loci that were differentially methylated with respect to temperature across tissues using a Fisher’s exact test. The distribution of temperature-associated DMCs with respect to gene context was also compared across tissues using Fisher’s exact tests followed by FDR correction for multiple comparisons. In this analysis, the representation of DMCs in each gene context was compared to the representation of all shared cytosines (N = 181,834 – (# DMCs)).

## Results

### Genome-wide DNA methylation patterns harbor signatures of hatchling sex and past incubation temperature

A total of 462,236 cytosines was retained in our analyses following filtering (denoted “covered cytosines” or “covered CpGs”). Covered CpGs make up a relatively small proportion (2.2%) of the 21,480,261 CG dinucleotides in the alligator genome. However, as has been previously demonstrated with RRBS techniques, covered CpGs were enriched in regions of potential functional significance relative to all the CpGs in the genome (Appendix S1; Figure S1), including in promoters (*P* < 0.0001; Figure S1A) and exons (*P* < 0.0001; Figure S1A), as well as CpG islands (*P* < 0.0001), CpG shores (*P* < 0.0001), and CpG shelves (*P* < 0.0001). Of the CpGs analyzed, 120 (0.026%) loci displayed sex-associated differential methylation patterns across all females and males irrespective of temperature (Figure 2; denoted “universal sex-associated DMCs”; Appendix S1). Similar proportions of these DMCs displayed greater methylation in females (n = 63; 52.5%) and greater methylation in males (n = 57; 47.5%; Figure 2A), with loci generally hypermethylated (> 75%) in one sex and exhibiting intermediate methylation in the other sex (Figure 2A, B). The mean difference in percent methylation between females and males was 23.8% for loci more highly methylated in females and 21.8% for loci more highly methylated in males (Figure 2C). Further, universal sex-associated DMCs were enriched in highly methylated loci (80-100% methylation in all samples; *P* = 0.039; Figure S1C). When we relaxed the requirement that loci be differentially methylated across all females and males, we identified 1,307 (0.28%) sex-associated DMCs. This group included loci that were only differentially methylated between sexes in hatchlings incubated at 34.5°C. Interestingly, only 10 CpGs were identified as differentially methylated across all females and males as well as within 34.5°C individuals, suggesting that a large proportion of sex-associated DMCs (n = 1,187; 90.8%) are specific to hatchlings incubated at 34.5°C.

**Figure 2.**
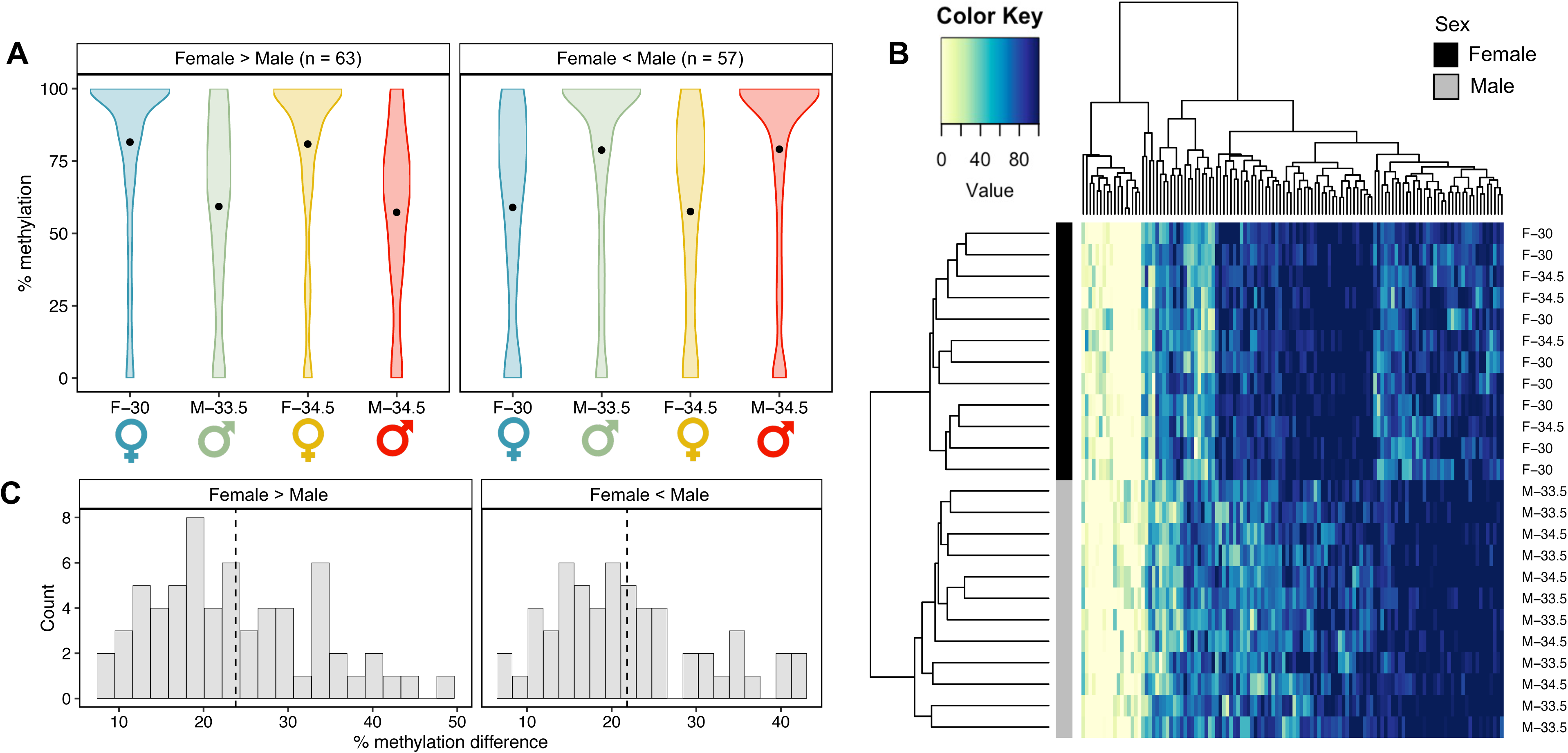
Sex-associated differential methylation patterns. (A) Violin plots depicting kernel density of the percent methylation of all universal sex-associated differentially methylated cytosines (DMCs; n = 120) for each treatment group, with panels separating DMCs that are hypermethylated in females (female-biased) from those that are hypermethylated in males (male- biased). Central points of violins represent overall mean percent methylation. (B) Heatmap depicting the percent methylation of all universal sex-associated DMCs across all samples. Each row represents a color-coded individual (black = female; grey = male), and each column represents a single DMC. Individual dendrogram positions were determined via hierarchical clustering. (C) Histograms of the difference in percent methylation for each DMC, with panels separating female-biased and male-biased DMCs. Vertical dotted line indicates mean difference in percent methylation.

Of the CpGs analyzed, 707 (0.15%) loci exhibited differential methylation with respect to incubation temperature and not sex (Figure 3). Similar proportions of these temperature-associated DMCs showed greater methylation with increasing temperatures (n = 356; 50.4%) and greater methylation with decreasing temperatures (n = 351; 49.6%). The mean difference in percent methylation between incubation temperature groups at these DMCs was 29.9% for loci that exhibited increased methylation with increased temperature and 29.5% for loci that exhibited decreased methylation with increased temperature (Figure 3C). Similar to universal sex-associated DMCs, temperature-associated DMCs tended to be highly methylated and were enriched in loci with an overall mean percent methylation between 80-100% relative to covered CpGs (*P* < 0.0001; Figure S1C). The comparison of 30°C hatchlings to 34.5°C yielded the greatest number of temperature-associated DMCs (n = 273), while the comparison of 30°C hatchlings to 33.5°C hatchlings yielded the least (n = 230). Few loci were identified as differentially methylated with respect to temperature across multiple pairwise temperature comparisons, and none were identified as significant across all three temperature comparisons. When testing for Spearman correlations between temperature and methylation status across all covered CpGs, we did not identify any significant correlations after FDR correction (Figure 4). Even so, several loci showed robust correlation coefficients with temperature (absolute correlation coefficients of ∼0.8; top 8 most correlated loci shown in Figure 4C). The absolute correlation coefficients of DMCs identified as temperature-associated through the pairwise comparisons were significantly greater than those loci not identified as temperature associated (*P* < 0.0001; Figure 4B).

**Figure 3.**
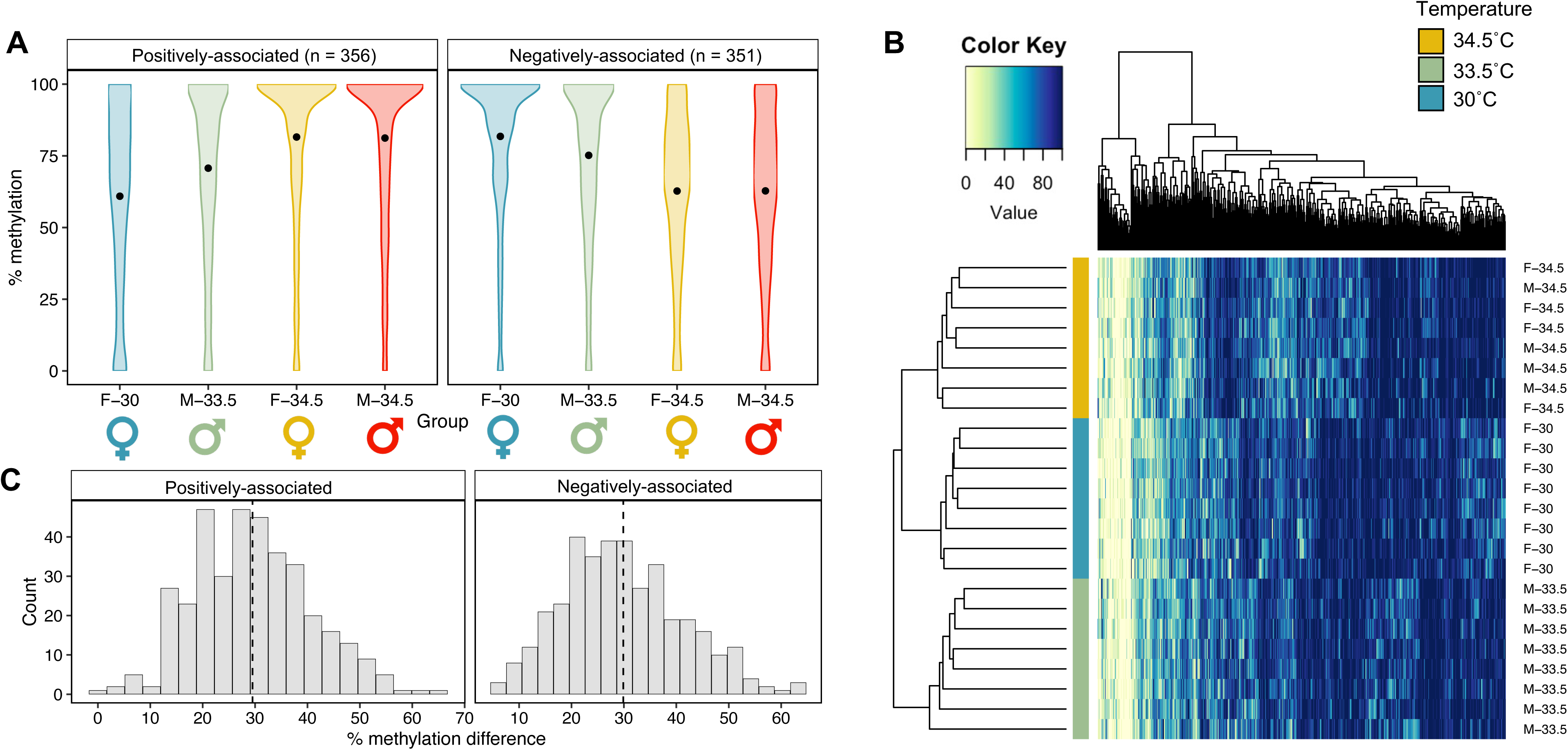
Temperature-associated differential methylation patterns. (A) Violin plots depicting kernel density of the percent methylation of all temperature-associated differentially methylated cytosines (DMCs; n = 707) for each treatment group, with panels separating DMCs that exhibit increased methylation with increasing temperature (positive) from those that exhibit decreased methylation with increasing temperature (negative). Central points of violins represent overall mean percent methylation. (B) Heatmap depicting the percent methylation of all temperature- associated DMCs across all samples. Each row represents a color-coded individual (yellow = 34.5°C; green = 33.5°C, blue = 30°C), and each column represents a single DMC. Individual dendrogram positions were determined via hierarchical clustering. (C) Histograms of the difference in percent methylation for each DMC, with panels separating positive temperature- associated DMCs from negative temperature-associated DMCs. Vertical dotted line indicates mean difference in percent methylation.

**Figure 4.**
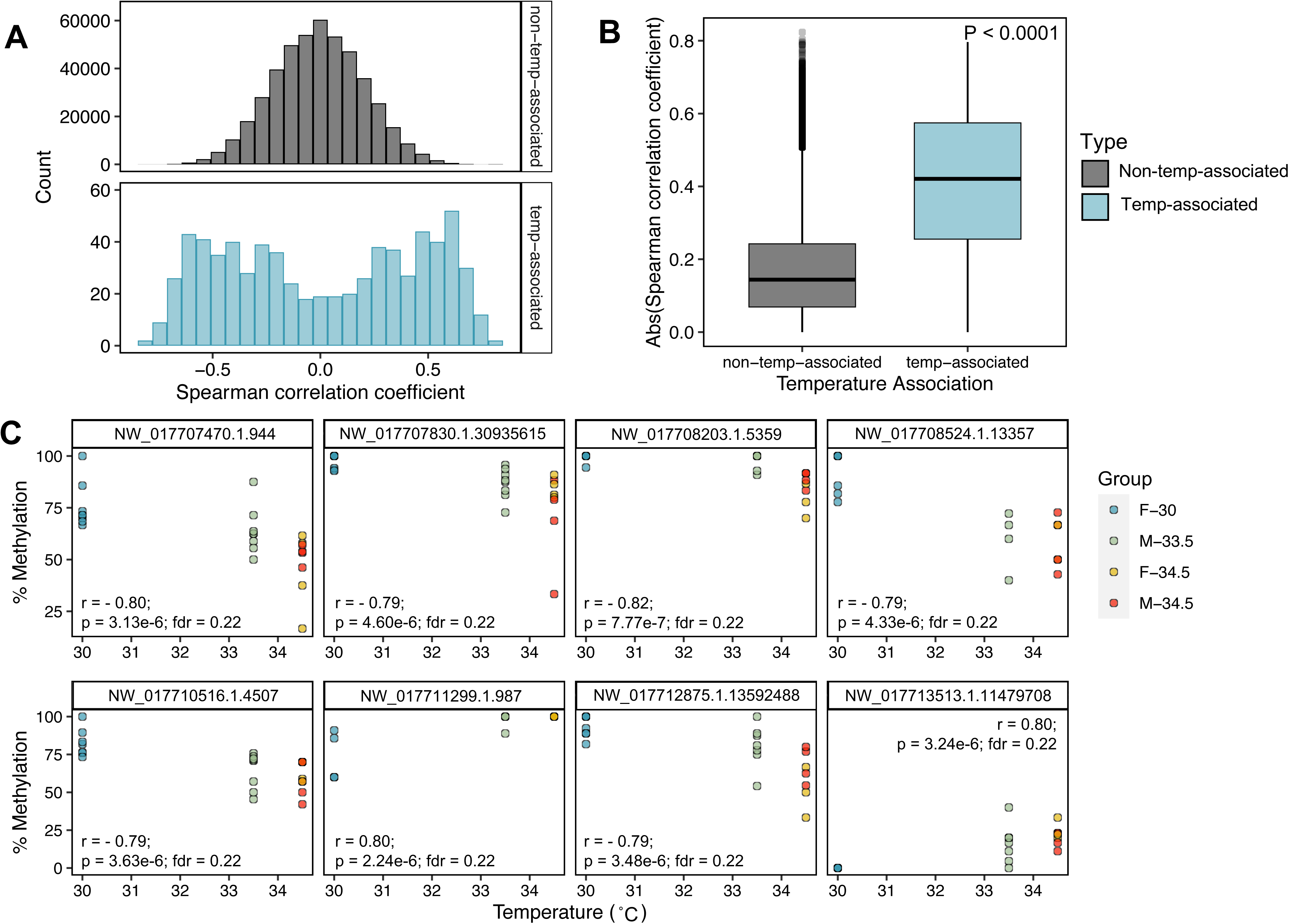
Correlations between incubation temperature and genome-wide DNA methylation patterns. (A) Histograms of Spearman correlation coefficients for temperature correlations, with panels separating loci not identified as temperature-associated via pairwise comparisons (top) from loci identified as temperature-associated differentially methylated cytosines (DMCs; bottom). (B) Boxplot comparing absolute Spearman correlation coefficients between temperature associated DMCs and all other loci. Central line of boxplot depicts median, box depicts interquartile range (IQR), vertical lines depict the maximum and minimum values, and values greater or less than 1.5*IQR are depicted as points. (C) Scatterplots depicting the relationship between percent methylation and incubation temperature for the loci exhibiting the top 8 most significant correlations with temperature.

### Differentially methylated cytosines do not show enrichment in promoters, but those that do overlap promoters and gene bodies tend to associate with genes involved in blood- and immune related processes

If the DMCs we identified are involved in gene regulation, we might expect them to be enriched in genomic regions of functional relevance such as promoters or gene bodies. We did not, however, observe such enrichment. In fact, sex-associated DMCs were significantly depleted in promoter regions (*P* = 0.00048) relative to covered CpGs and enriched in intergenic regions (*P* = 0.013; Figure S1A), but this pattern was not observed when only considering the universal sex-associated DMCs identified across all hatchlings. These DMCs, as well as temperature-associated DMCs, exhibited a similar distribution across different gene contexts to all the covered CpGs (Figure S1A), with subsets of DMCs overlapping with promoters (9.2% of universal sex-associated DMCs, 6.5% of sex-associated DMCs, and 7.6% of temperature associated DMCs) and gene bodies (45.8% of universal sex-associated DMCs, 39.3% of sex-associated DMCs, and 41.6% of temperature associated DMCs). Genes with sex-associated DMCs within their promoters were enriched for the GO term “DNA binding domain binding” (GO:0050692) (Table S3), and genes with temperature-associated DMCs within their promoters were enriched for the GO terms “integral component of synaptic membrane” (GO:0099699), “cell-cell contact zone” (GO: 0044291), “early phagosome” (GO:0032009), and “intrinsic component of synaptic membrane” (GO:0099240) (Table S3). When we considered genes with DMCs within their promoters or gene bodies, we identified several more enriched GO terms, with the blood-related GO terms “vasculature development” (GO:0001944) and “blood vessel development” (GO:0001568) enriched for genes with temperature-associated DMCs within their promoters or gene bodies (Table S4).

### Sex-associated differentially methylated cytosines collectively exhibit correlations with estradiol concentrations

Consistent with the results of (McCoy et al., 2016a), we did not detect a significant effect of sex or incubation temperature on plasma E2 concentrations, though the effect of sex was close to the threshold for significance (α = 0.05; *F*_1,18.91_ = 3.87; *P* = 0.064; Figure 5A). We also did not detect significant Spearman correlations between plasma E2 and the methylation status of any one cytosine after the FDR correction, though several loci showed robust correlation coefficients (top 8 most correlated loci shown in Figure 5D). Sex-associated DMCs did, however, show significantly greater absolute correlation coefficients with E2 concentrations than loci that were not sex-associated (Figure 5B, C). The same was not observed for T concentrations (Figure S3). The apparent link to estrogen signaling was not reflected in the proximity of sex-associated DMCs to putative EREs, nor AREs, as this did not differ significantly from that of covered CpGs (Figure S2,4).

**Figure 5.**
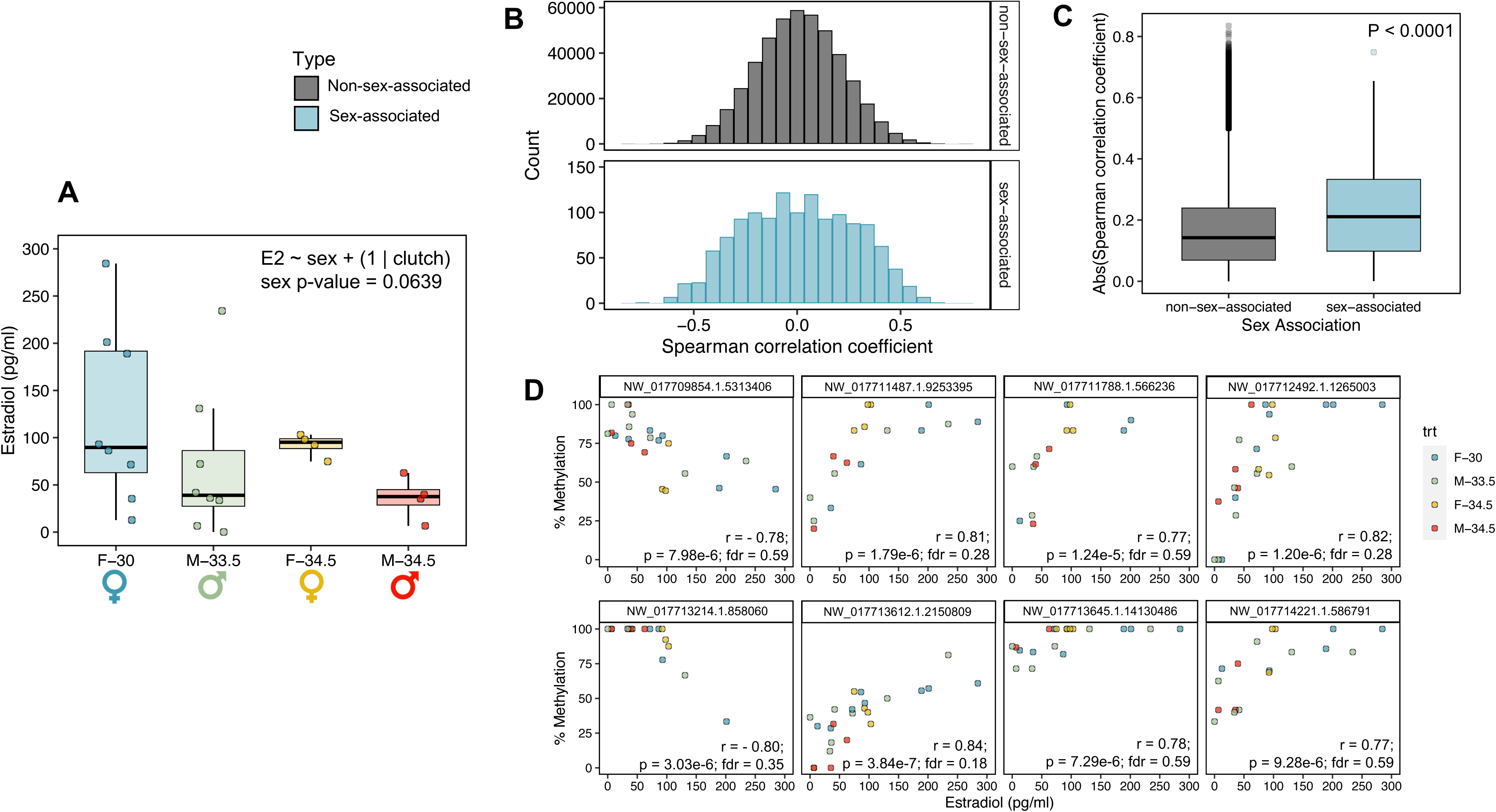
Correlations between plasma estradiol (E2) concentrations and genome-wide DNA methylation patterns. (A) Boxplot comparing plasma E2 concentrations between treatment groups. (B) Histograms of Spearman correlation coefficients for E2 correlations, with panels separating loci not identified as sex-associated via pairwise comparisons (top) from loci identified as sex-associated differentially methylated cytosines (DMCs; bottom). (C) Boxplot comparing absolute Spearman correlation coefficients between sex-associated DMCs and all other loci. (C) Scatterplots depicting the relationship between percent methylation and plasma E2 concentration for the loci exhibiting the top 8 most significant correlations. For boxplots, central line depicts median, box depicts interquartile range (IQR), vertical lines depict the maximum and minimum values, and values greater or less than 1.5*IQR are depicted as points.

### Methylation patterns of separate subsets of cytosines predict hatchling sex and past incubation temperature

Beyond describing sex- and temperature-associated DNAm patterns, we also aimed to determine whether DNAm patterns in blood cells can be used to predict the sex and/or past incubation temperature of hatchlings for which this information is unknown. We developed DNAm-based predictors of sex and past incubation temperature based on the methylation status of a subset of loci. For the predictive model of hatchling sex, 23 cytosines passed the threshold to be included in variable selection via elastic net regularization, and 20 of those cytosines were selected to be included in the final model (Table S5). This model correctly assigned sex to all 20 of the training set individuals with minimal variability in the model predictions (Mean ± SE predicted probability male: Females = 0.07 ± 0.01; Males = 0.93 ± 0.01; Figure 6A). This model also correctly assigned sex to all 4 of the test set individuals, though there was more variability in the model predictions (Mean ± SE predicted probability male: Females = 0.28 ± 0.07; Males = 0.71 ± 0.01; Figure 6B). For the predictive model of past incubation temperature, 152 cytosines passed the threshold to be included in variable selection, and 24 of those cytosines were selected to be included in the final model (Table S6). This model fit the training set well, with an R-squared greater than 0.99 between the actual incubation temperatures and fitted incubation temperatures. The mean absolute error for the training set individuals was 0.05°C (Figure 6C). However, the model did not perform as well on the test set individuals. The R-squared for the test set was 0.81 and the mean absolute error of the model predictions was 1.20°C (Figure 6D).

**Figure 6.**
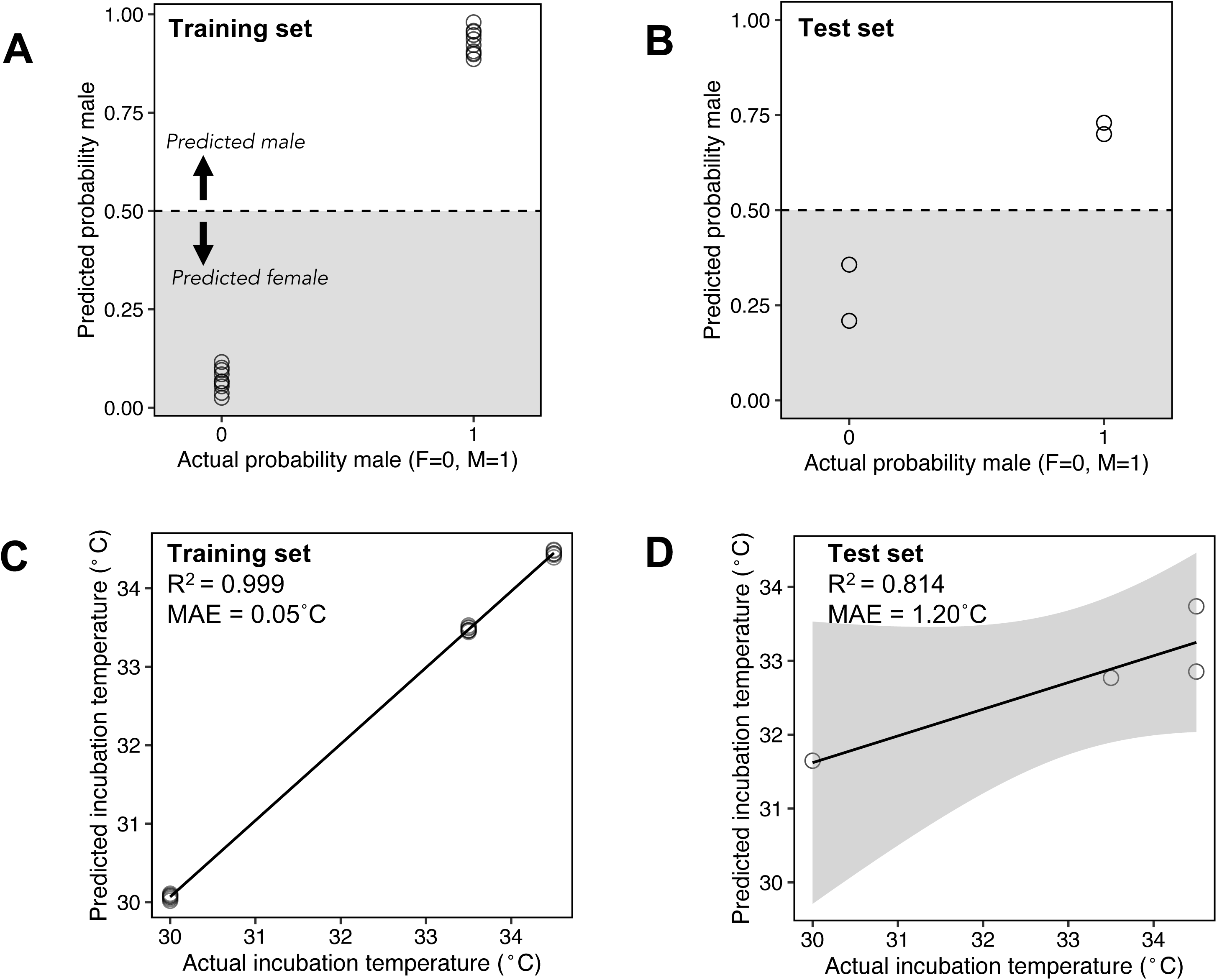
Predictive models of hatchling sex and past incubation temperature based on the methylation status of a subset of cytosines. (A) Hatchling sex predictions resulting from penalized binomial logistic regression with elastic net regularization for training set (n = 20). (B) Hatchling sex predictions for test set (n = 4). (C) Incubation temperature predictions resulting from penalized linear regression with elastic net regularization for training set (n = 20). (D) Incubation temperature predictions for test set (n = 4). The model predicting hatchling sex was based on the methylation status of 20 loci, while the model predicting incubation temperature was based on the methylation status of 24 loci. MAE = mean absolute error.

### Embryonic gonads following temperature-dependent sex determination show more extensive temperature-associated DNA methylation patterning compared to hatchling blood cells

Though we observed both sex- and temperature-associated patterns in the DNA methylomes of hatchling blood cells, these patterns were considerably less extensive than the temperature-associated DNAm patterns we observed in the embryonic gonad following TSD (Figure 7). In our comparison of the blood and gonadal tissues, we used information on 181,834 CpG sites (39.3% of covered CpGs in blood) which overlapped both RRBS datasets. Of these shared, covered cytosines, 232 (0.13%) were identified as differentially methylated with respect to incubation temperature in blood, whereas 4,339 (2.39%) cytosines were identified as differentially methylated with respect to incubation temperature in the gonad (Figure 7A). The proportion of DMCs identified in the gonad was significantly greater than that identified in blood (*P* < 0.0001). Only five loci were identified as temperature-associated DMCs in both tissues and the direction of the methylation difference was not consistent across tissues. The maximum absolute difference in percent methylation for DMCs identified in the gonad was 100%, while the maximum absolute different in percent methylation for DMCs identified in the blood was 66.67% (Figure 7B). Temperature-associated DMCs identified in the blood displayed a similar genomic distribution with respect to gene context as all shared, covered CpGs (Figure 7C). However, temperature-associated DMCs in the gonad were depleted in promoters and enriched in introns relative to all shared, covered CpGs (Figure 7C).

**Figure 7.**
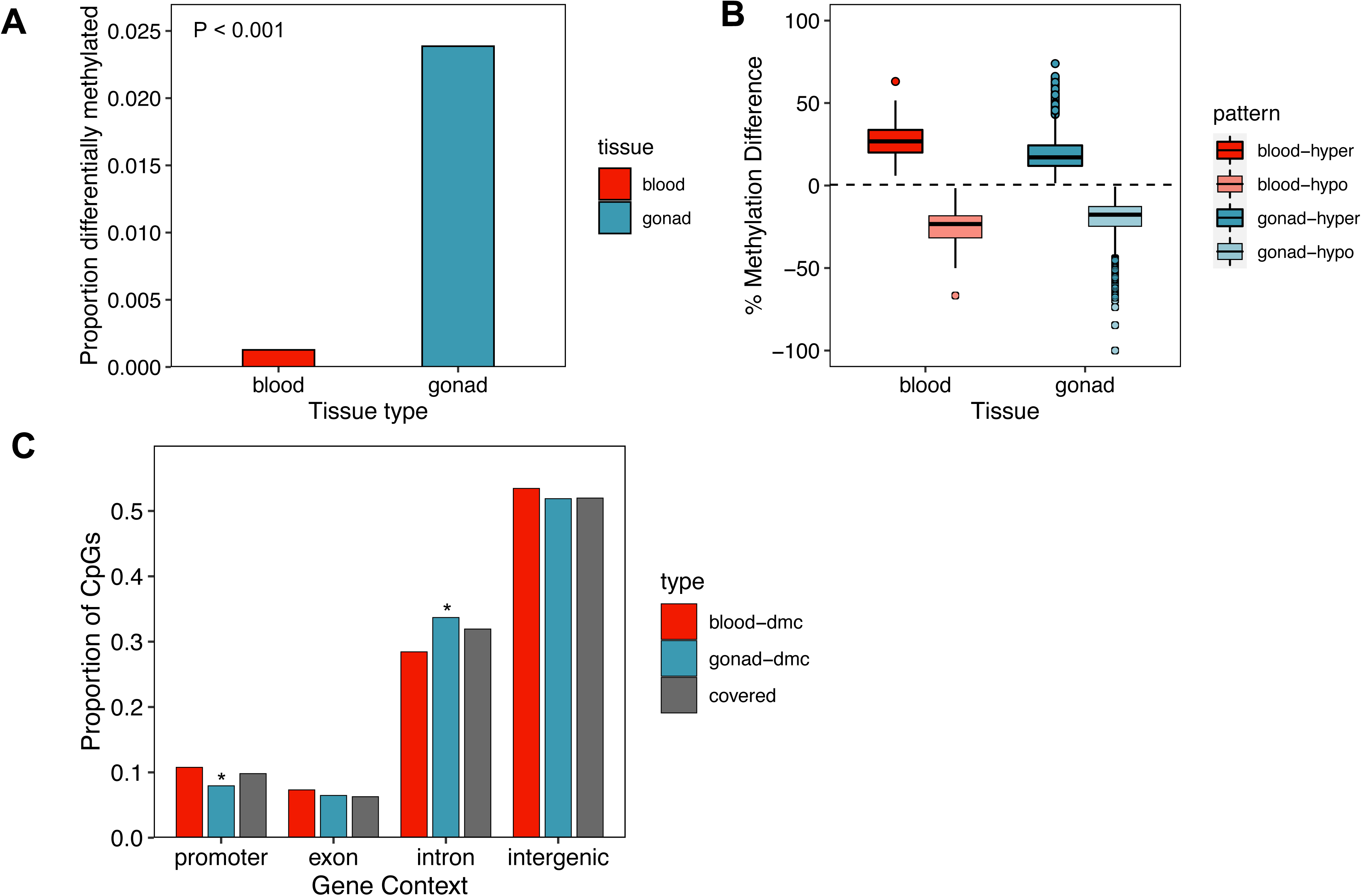
Comparison of temperature-associated DNA methylation patterns in blood versus embryonic gonad just after temperature-dependent sex determination. (A) Proportion of covered CpG sites shared among blood and gonad RRBS data (n =181,834) that are differentially methylated with respect to incubation temperature. P-value is the derived from a Fisher’s exact test to determine if the proportion of differentially methylated versus non-differentially methylated loci is different between tissues. (B) Comparison of percent methylation difference (i.e., effect size) of temperature-associated differentially methylated cytosines (DMCs) identified in each tissue. (C) Comparison of the distribution of temperature-associated DMCs identified in each tissue with respect to gene context. Asterisk indicates significant difference in proportion of DMCs in a context category compared to the shared covered cytosines at an FDR threshold of 10%. The only significant differences we observed was depletion of gonadal temperature-associated DMCs in promoter regions and enrichment of gonadal temperature-associated DMCs in introns.

## Discussion

In the present study, we demonstrate that the DNA methylomes of blood cells from hatchling alligators are sexually dimorphic and retain signatures of past incubation temperature. We further show that DNAm patterns at a subset of loci can be used to predict sex and past incubation temperature. It remains unclear whether the observed sex- and temperature-associated DNAm patterns are functionally linked to the transcriptional activity of blood cells or are passive, indirect markers of functional changes occurring in other tissues. Like most non-mammalian vertebrates, reptiles possess nucleated red blood cells that are transcriptionally active (Waits et al., 2020). The observed link between DMCs and genes with blood- and immune-related gene ontology terms suggests these DNAm patterns may, at least in part, be connected to gene regulatory responses in blood cells. It should be noted, however, that information on blood cell composition was not incorporated into the present study and may contribute to the observed DNAm patterns (Lea et al., 2017), though we do not have prior evidence to suggest blood cell composition would differ across treatment groups. Collectively, our results raise intriguing questions regarding tissue-specific epigenomic patterning in the context of developmental plasticity and support the potential utility of DNAm-based markers as tools for ecological monitoring and conservation in the face of environmental change. Further, these findings point to several critical considerations relevant to the implementation of epigenomic tools in conservation more broadly.

During TSD, the gonadal DNA methylome undergoes widespread remodeling in response to incubation temperature. These epigenomic changes likely play roles in both the establishment of sexual fate during the discrete window of development in which sex determination is sensitive to temperature as well as the maintenance of sexual fate into adulthood (Navarro-Martín et al., 2011; Parrott, Kohno, et al., 2014; Weber & Capel, 2021). Compared to temperature-associated DNAm patterns in embryonic gonads immediately following sex determination, temperature influences on the DNA methylomes of hatchling blood cells were considerably less extensive. Further, the loci acquiring temperature-associated DNAm patterns appeared largely unique to each tissue. This is perhaps not surprising considering DNAm patterns play a key role in the maintenance of cellular identity (Bock et al., 2012; Ziller et al., 2013) and thus tend to diverge markedly depending on tissue and cell type (Blake et al., 2020; Christensen et al., 2009). Nonetheless, previous studies have identified subsets of loci displaying consistent DNAm responses to environmental cues across tissues, termed ‘metastable epialleles’(Bourc’his, 2021; Rakyan et al., 2002). In a study of European sea bass, nine genes showed consistent differential methylation related to past developmental temperature across all tissues examined, including brain, muscle, liver, and testis (Anastasiadi et al., 2021). While results from the present study suggest DNAm responses to incubation temperature are largely tissue-specific, it should be noted that our interpretation is limited by the fact that the blood and gonadal samples used in this study were derived from independent sets of individuals. Further, in comparing tissues, we limited our analyses to temperature-associated DNAm patterns because sexual fate was unknown in the embryonic samples. By examining the DNA methylome across different tissues within the same individuals and examining sex-associated epigenetic patterns in addition to those associated with temperature, future studies are likely to yield valuable information regarding the prevalence and biological significance of metastable epialleles.

The observation that DNA methylomes of blood cells retain signatures of past incubation temperature in hatchlings reared in a common post-natal environment raises several interesting questions regarding the persistence of environmentally induced DNAm patterns established during development. In particular, how does the plasticity of DNAm vary across ontogeny and across tissues? DNAm patterns tend to be more dynamic early in life and lasting impacts of environmental cues experienced during development on the epigenome have been observed across diverse contexts (Angers et al., 2010; Barua et al., 2014; Borghol et al., 2012; Dolinoy et al., 2007; Heijmans et al., 2008). Even within the embryonic period, certain developmental windows are characterized by unique environmental sensitivity (Faulk & Dolinoy, 2011). In the context of TSD, the thermal plasticity of gonadal fates is limited to a specific period of development termed the thermosensitive period during which sexually dimorphic epigenomic patterns are established in response to temperature (Ge et al., 2018; McCoy et al., 2015). These gonadal DNAm patterns are then maintained across mitotic cell divisions. However, somatic tissues are likely to differ in their pattern of epigenetic thermal plasticity based on multiple factors including cell type-specific transcriptional activity (Christensen et al., 2009). While hatchling blood cells exhibited temperature-associated DNAm patterns that persisted until at least seven days post-hatch, it is unclear whether these patterns remain in juveniles and adults. Further, if hatchlings experienced different post-natal environments, would variation in current thermal cues override the influences of incubation temperature on DNAm patterns? How current and past environmental signals are integrated at the level of the DNA methylome remains largely unknown (Lea et al., 2016; Metzger & Schulte, 2017). Yet this represents an important topic of future inquiry from both basic and applied perspectives.

If signatures of past incubation temperature indeed persist in the DNA methylomes of adult blood cells, these patterns could be applied to address diverse ecological and evolutionary questions that are currently inaccessible to studies of long-lived taxa. For instance, current demographic structure of wild populations could be linked to past incubation conditions to reveal the relationships between historical environmental heterogeneity and contemporary population-level variation. Further, one of the primarily obstacles to understanding the evolutionary significance of developmental plasticity in reptiles is the inability to follow the fitness trajectories of individuals from incubation to reproductive maturity, which often occurs decades later (Mitchell et al., 2018). The ability to retroactively determine incubation temperature in adults would undoubtedly transform research of this nature. However, beyond examining the degree to which epigenetic signatures of incubation temperature persist in later life stages, future studies must address a few other critical questions. To what extent do DNAm patterns reflect fluctuations in developmental thermal cues? Wild nests exhibit variable thermal regimes (Bock et al., 2020; Carter et al., 2018; Escobedo-Galván et al., 2016; Warner & Shine, 2011), thus applications of DNAm markers to understand incubation conditions in the field must account for the role of thermal fluctuations in influencing these patterns. Further, the present study was limited in sample size and number of temperature treatments. Future studies including a broader range of incubation temperatures as well as a larger model training set are likely to yield a DNAm predictor capable of estimating past incubation temperature with greater accuracy.

One of the aims of this study was to identify loci for which DNAm was predictive of sex and thus could be used as a non-lethal method of sexing TSD species in early-life stages. We identified 20 cytosines in the alligator genome for which methylation status was predictive of phenotypic sex. Further, 120 cytosines exhibited sex-associated variation in methylation entirely independent of incubation temperature. While female and male hatchlings did not differ significantly in plasma E2 concentrations, sex-associated DMCs, as a whole, exhibited greater correlations between E2 and methylation status than those not associated with sex. This suggests the DNA methylome may reflect more subtle estrogen-directed responses that are not apparent at the level of plasma concentrations, consistent with previous work linking DNAm patterns to endogenous sex hormone signaling and exposure to estrogenic contaminants (Almstrup et al., 2016; Hanna et al., 2012; Nugent et al., 2015; Zhao et al., 2017). Overall, sex-associated DNAm patterns in blood cells represent promising candidates to serve as non-lethal indicators of phenotypic sex in early-life stages of TSD species. However, genome-wide approaches to profiling DNAm are not currently a practical option for conservation efforts to monitor hatchling sex ratios in the field. The utility of these DNAm marks as a conservation tool depends on validation using more targeted approaches (e.g., bisulfite sequencing PCR) across many samples originating from multiple geographic sites. Further, a quantitative comparison of the reliability and cost-efficiency of different potential molecular markers of sex (e.g., blood DNAm patterns, sex-biased protein levels (Tezak et al., 2020), high-sensitivity steroid hormone quantitation (Bae et al., 2021)) in TSD species is warranted.

In the face of rapidly changing environmental conditions, monitoring the responses of key demographic parameters, like offspring sex ratios, to climate change in real time will play a critical role in the conservation of thermally sensitive species. This is the first study to use somatic DNAm patterns to predict phenotypic sex in a species with environmental sex determination. Intriguingly, we also demonstrate that DNAm patterns hold the potential to provide information about past incubation temperatures, though this finding requires validation in the context of fluctuating incubation temperatures as well as in older animals. Understanding how genetic and environmental variation contribute to these methylation patterns represents a critical target for future research that will advance efforts to realize the full potential of epigenomic tools in conservation contexts.

## Acknowledgements

We thank past and present members of the Parrott Lab for their significant contributions to egg incubation experiments and helpful discussions regarding the analysis and interpretation of our findings. We are grateful to Dr. Louis Guillette Jr. and past members of his research group for their contributions to the initial hatchling experiment, without which this work would not have been possible. We also thank Arnold Brunell and other colleagues from the Florida Fish and Wildlife Commission for their assistance with permitting and field support during egg collections. This work was supported in part by a grant from the IUCN Crocodile Specialist Group Student Research Assistance Scheme (to SLB), a grant from the Joshua Laerm Academic Support Fund of the Georgia Museum of Natural History (to SLB), and a grant from the National Science Foundation (Award # 1754903 to BBP). Furthermore, this material is based upon work supported by the Department of Energy Office of Environmental Management (Award # DE- EM0005228 to the University of Georgia Research Foundation).

## Disclaimer

This report was prepared as an account of work sponsored by an agency of the United States Government. Neither the United States Government nor any agency thereof, nor any of their employees, makes any warranty, express or implied, or assumes any legal liability or responsibility for the accuracy, completeness, or usefulness of any information, apparatus, product, or process disclosed, or represents that its use would not infringe privately owned rights. Reference herein to any specific commercial product, process, or service by trade name, trademark, manufacturer, or otherwise does not necessarily constitute or imply its endorsement, recommendation, or favoring by the United States Government or any agency thereof. The views and opinions of authors expressed herein do not necessarily state or reflect those of the United States Government or any agency thereof.

## Data Availability Statement

Upon acceptance, raw sequencing reads and associated metadata from reduced-representation bisulfite sequencing libraries will be deposited in the NCBI Sequence Read Archive (SRA), additional related data will be available on Dryad, and scripts used in data analysis will be available on Github.

## Author Contributions

S.L.B. and B.B.P. conceived of the study. J.A.M., S.L.B., and B.B.P. designed and conducted incubation experiments and collected samples. S.L.B. performed nucleic acid extraction and sequencing library preparation. S.L.B. performed data analysis. C.R.S. and B.B.P. assisted with data analysis. S.L.B. and B.B.P. drafted the initial manuscript which was subsequently edited by all authors.

