## Supplemental Information for "Genome-wide DNA methylation patterns harbor signatures of hatchling sex and past incubation temperature in a species with environmental sex determination"

**Supplemental Information**
**Table of Contents**


| **Content** | **Pages** |
|---|---|
| **Appendix S1.** Supplementary Materials and Methods. | **1** |
| **Table S1.** Incubation experiments to characterize the American alligator (*Alligator mississippiensis*) temperature-by-sex ratio reaction norm. | **2-4** |
| **Table S2.** Sequencing summary for reduced-representation sequencing libraries from hatchling blood cells. | **5** |
| **Table S3.** Gene ontology analysis for genes associated with differentially methylated cytosines overlapping promoter regions. | **6** |
| **Table S4.** Gene ontology analysis for genes associated with differentially methylated cytosines overlapping promoter regions or gene bodies. | **7-11** |
| **Table S5.** Model coefficients for cytosines selected to be included in the predictive model of sex. | **12** |
| **Table S6.** Model coefficients for the cytosines selected to be included in the predictive model of incubation temperature. | **13-16** |
| **Table S7.** Sequencing summary for reduced-representation sequencing libraries from embryonic gonads. | **17** |
| **Figure S1.** Genomic characterization of sex- and temperature-associated differentially methylated cytosines with respect to gene context and CpG density. | **18** |
| **Figure S2.** Associations between differentially methylated loci and putative estrogen-response elements. | **19** |
| **Figure S3.** Correlations between plasma testosterone concentrations and genome-wide DNA methylation patterns. | **20** |
| **Figure S4.** Associations between differentially methylated cytosines and putative androgen response elements. | **21** |

**Supplemental Information**
Appendix S1. Supplementary Materials and Methods.

*Detailed protocol for blood nucleic acid isolation and quantification*

DNA was isolated from blood cells using a modified column approach based on the SV total RNA isolation system (Promega). Frozen samples were thawed on ice. To remove RNAlater from the preserved cells, 500 µl of phosphate-buffered saline (PBS) was added to a 200 µl aliquot of each sample and samples were centrifuged at 5000 rcf for 15 minutes at 4˚C (and for an additional 5 min if a cell pellet did not form). Pelleted cells were washed once more with 500 µl of PBS and nucleic acids were isolated according to the protocol reported in (Bae et al., 2021). Specifically, 350 µl of lysis buffer (4 M guanidinium thiocyanate; 0.01 M Tris-HCl, pH 7.5; 2% beta-mercaptoethanol) was added to each sample, and samples were homogenized with a sterile steel bead using a Mini-BeadBeater 24 (BioSpec) at 30 Hz for 2 min. Homogenized samples were then centrifuged for 3 min at 14,000 rcf, and the resulting supernatants transferred to spin columns with silica membranes (Epoch Life Science) and centrifuged for 30 sec at 14,000 rcf to bind DNA to the column. Columns were then washed twice with 700 µl wash buffer (162.8 mM potassium acetate; 27.1 mM Tris-HCl, pH 7.5; diluted with 60% (v/v) ethanol) and DNA was eluted in 60 µl TE buffer (pH 8.0).

*Reduced-representation bisulfite sequencing enriches for genomic regions of potential functional significance*

A total of 462,236 cytosines was retained in our analyses following filtering, which are referred to as "covered cytosines" or "covered CpGs". Covered CpGs make up a relatively small proportion (2.2%) of the 21,480,261 CG dinucleotides in the alligator genome. However, as has been previously demonstrated with RRBS techniques, *MspI* digestion followed by size selection resulted in covered CpGs that were enriched in regions of potential functional significance relative to all the CpGs in the genome, including in promoters ($P < 0.0001$; **Figure S1A**) and exons ($P < 0.0001$; **Figure S1A**), as well as CpG islands ($P < 0.0001$), CpG shores ($P < 0.0001$), and CpG shelves ($P < 0.0001$). Covered CpGs were also enriched in intergenic regions ($P < 0.0001$; **Figure S1A**) relative to all CpGs and depleted in introns ($P < 0.0001$; **Figure S1A**) and open sea regions ($P < 0.0001$). Consistent with their distribution with respect to CpG island context, covered CpGs were depleted in genomic tiles of low CpG density (i.e., those containing between 0 and 20 CpGs per 1 kb) and were enriched in genomic tiles of intermediate and high CpG densities (i.e., those containing 20 to 40, 40 to 60, and 60 to 80 CpGs per 1 kb; **Figure S1B**). Covered CpGs were also enriched in genomic regions in proximity to putative EREs ($P < 0.0001$; **Figure S2**) relative to all CpGs in the genome, but not in genomic regions in proximity to putative AREs (**Figure S4**).

*Definitions of sex-associated differentially methylated cytosines.*

Cytosines that were identified as differentially methylated between all females and males (FvM) *or* between 34.5˚C females and 34.5˚C males ($F^{34.5}vM^{34.5}$) are collectively referred to as "sex-associated DMCs." If referring to *only* those DMCs identified between all females and males, the term "universal sex-associated DMCs" is used.

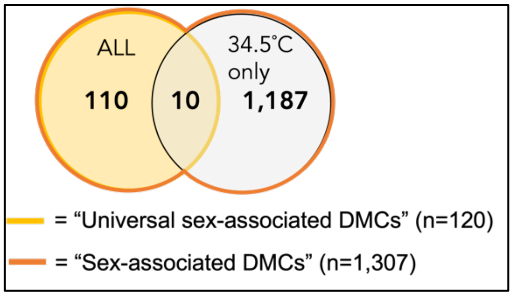

Table S1. Incubation experiments to characterize the American alligator (*Alligator mississippiensis*) temperature-by-sex ratio reaction norm.

| Study ID | Reference | Incubation Temperature | # female | # male | % female | Method of sexing | Egg collection and other notes |
|---|---|---|---|---|---|---|---|
| A | (Joanen, McNease, & Ferguson, 1987) | 32.8 | 0 | 106 | 0 | | |
| A | | 31.7 | 13 | 38 | 0.25490196 | | |
| A | | 30.6 | 19 | 13 | 0.59375 | | |
| A | | 29.4 | 80 | 0 | 1 | | |
| B | (Janes et al., 2013) | 28 | 10 | 0 | 1 | Macroscopic inspection (Moore et al., 2008) | Field collection, "late-stage embryos" |
| B | | 33 | 0 | 10 | 0 | | |
| C | (Lance & Bogart, 1991) | 30 | 25 | 0 | 1 | Histology | Eggs collected from wild nests within 1-2 days of oviposition; subset injection of vehicle (corn oil) |
| C | | 33 | 0 | 27 | 0 | | |
| D | (Lance & Bogart, 1992) | 30 | 58 | 0 | 1 | Histology | Subset injected with vegetable oil |
| D | | 33 | 0 | 56 | 0 | | |
| E | (Lang & Andrews, 1994) | 29 | 113 | 0 | 1 | Macroscopic inspection of gonads or genitalia at various ages | Eggs collected from wild nests within 1 week of oviposition |
| E | | 30 | 27 | 0 | 1 | | |
| E | | 30.5 | 27 | 0 | 1 | | |
| E | | 31 | 197 | 0 | 1 | | |
| E | | 31.5 | 125 | 0 | 1 | | |
| E | | 32 | 114 | 228 | 0.33333333 | | |
| E | | 32.5 | 0 | 122 | 0 | | |
| E | | 33 | 0 | 97 | 0 | | |
| E | | 33.5 | 3 | 16 | 0.15789474 | | |
| E | | 34 | 35 | 19 | 0.64814815 | | |
| E | | 34.5 | 14 | 1 | 0.93333333 | | |
| F | (Milnes et al., 2004) | 32 | 11.62 | 2.38 | 0.83 | Histology | Eggs collected from wild nests within 2 weeks of oviposition; unsure of exact sample sizes, values reported are the max possible based on methods |
| F | | 33.5 | 0 | 14 | 0 | | |
| G | (Milnes et al., 2005) | 32 | 5 | 5 | 0.5 | Histology | Eggs collected from wild nests within 2 weeks of oviposition |
| G | | 33.5 | 0 | 10 | 0 | | |
| H | (Moore et al., 2010) | 30 | 12 | 0 | 1 | Macroscopic inspection of gonad, presence of oviduct at 13 months | Included eggs collected from Lake Woodruff and Lake Apopka; sex ratio also includes the effect of survival to 13 months |
| H | | 33.5 | 0 | 13 | 0 | | |
| H | | 32 | 8 | 19 | 0.2962963 | | |
| I | (Guillette Jr. et al., 1994) | 30.5 | 24 | 14 | 0.63157895 | Histology | Average age of egg at collection was 12 days; included individuals from both Lake Apopka and Lake Woodruff |
| J | (Conley et al., 1997) | 31 | 14 | 0 | 1 | Histology (between stage 25 and hatching) | Data included here are from 1995 study |
| J | | 33 | 0 | 10 | 0 | | |
| J | | 34 | 11 | 12 | 0.47826087 | | |
| J | | 34.2 | 5 | 0 | 1 | | |
| K | (Ferguson & Joanen, 1983) | 30 | 97 | 0 | 1 | Histology at day 60 of incubation | Eggs collected within 10hr of oviposition |
| K | | 32 | 85 | 13 | 0.86734694 | | |
| K | | 34 | 0 | 94 | 0 | | |
| L | (Deeming & Ferguson, 1991) | 30 | 19 | 0 | 1 | Macroscopic inspection of gonad at day 60 of incubation | Only included control individuals from each study year |
| L | | 33 | 0 | 19 | 0 | | |
| M | (Bock et al., 2021) | 30 | 68 | 1 | 0.98550725 | | Included all treatment groups |
| M | | 33.5 | 1 | 62 | 0.01587302 | | |

| Study ID | Reference | Incubation Temperature | # female | # male | % female | Method of sexing | Egg collection and other notes |
|---|---|---|---|---|---|---|---|
| M | | 31.2 | 95 | 0 | 1 | Macroscopic inspection of gonad, presence of oviduct at 10 days post-hatch | |
| N | (McCoy et al., 2016) | 30 | 19 | 0 | 1 | AMH and CYP19A1 expression | All eggs collected from wild nests at Lake Apopka, FL |
| N | | 32 | 41 | 15 | 0.73214286 | | |
| N | | 33.5 | 4 | 16 | 0.2 | | |
| N | | 34.5 | 12 | 46 | 0.20689655 | | |

Table S2. Sequencing summary for reduced-representation sequencing libraries from hatchling blood cells.

| Sample | Temperature | Sex | Clutch | Total reads | Total aligned reads | Percent alignment | Bisulfite conversion efficiency |
|---|---|---|---|---|---|---|---|
| 1 | 30 | F | AP-03 | 6525084 | 4213047 | 64.5669389 | 99.6197 |
| 2 | 33.5 | M | AP-06 | 10336198 | 6617520 | 64.0227674 | 99.3485 |
| 3 | 33.5 | M | AP-08 | 7501614 | 4905246 | 65.3892082 | 99.1967 |
| 4 | 34.5 | F | AP-06 | 10819612 | 6718596 | 62.0964597 | 99.5743 |
| 5 | 33.5 | M | AP-04 | 8676405 | 5430747 | 62.5921335 | 99.5053 |
| 6 | 34.5 | M | AP-06 | 8170904 | 5098412 | 62.3971595 | 99.7712 |
| 7 | 30 | F | AP-04 | 9064806 | 5734847 | 63.2649722 | 99.6243 |
| 8 | 34.5 | M | AP-03 | 9038702 | 5882088 | 65.0766891 | 99.3175 |
| 9 | 30 | F | AP-03 | 10732905 | 6762810 | 63.0100611 | 99.8536 |
| 10 | 30 | F | AP-06 | 9106305 | 5885739 | 64.6336687 | 99.896 |
| 11 | 33.5 | M | AP-04 | 7543166 | 4976851 | 65.9782776 | 99.3638 |
| 12 | 34.5 | M | AP-03 | 10512637 | 6545212 | 62.2604205 | 99.5369 |
| 13 | 33.5 | M | AP-03 | 7589151 | 4805508 | 63.3207588 | 99.6802 |
| 14 | 33.5 | M | AP-03 | 8129634 | 5234303 | 64.3854693 | 99.4774 |
| 15 | 30 | F | AP-08 | 8422219 | 5270330 | 62.5765015 | 99.4653 |
| 16 | 33.5 | M | AP-08 | 7724544 | 4807692 | 62.2391691 | 99.9292 |
| 17 | 34.5 | F | AP-03 | 8734654 | 5610949 | 64.2377935 | 99.8738 |
| 18 | 33.5 | M | AP-05 | 8310169 | 5387366 | 64.8285973 | 99.1258 |
| 19 | 30 | F | AP-05 | 9557288 | 6006117 | 62.8433192 | 99.889 |
| 20 | 30 | F | AP-08 | 8069098 | 5006391 | 62.043998 | 99.8269 |
| 21 | 34.5 | F | AP-04 | 7613576 | 4990415 | 65.5462689 | 99.3034 |
| 22 | 34.5 | F | AP-03 | 8080586 | 4994461 | 61.8081535 | 99.2036 |
| 23 | 30 | F | AP-04 | 7639275 | 4849357 | 63.479283 | 99.5237 |
| 24 | 34.5 | M | AP-04 | 8965500 | 5765134 | 64.3035414 | 99.7648 |

Table S3. Gene ontology analysis for genes associated with differentially methylated cytosines overlapping promoter regions.

| **Sex-associated DMCs** (across all or within 34.5˚C individuals) | | | |
|---|---|---|---|
| Source | GO term name | GO term ID | Benjamini-Hochberg FDR |
| GO:MF | DNA binding domain binding | GO:0050692 | 0.099478029 |
| **Universal sex-associated DMCs** (across all individuals) | | | |
| Source | GO term name | GO term ID | Benjamini-Hochberg FDR |
| NA | NA | NA | NA |
| **Temperature-associated DMCs** | | | |
| Source | GO term name | GO term ID | Benjamini-Hochberg FDR |
| GO:CC | integral component of synaptic membrane | GO:0099699 | 0.066156435 |
| GO:CC | cell-cell contact zone | GO:0044291 | 0.066156435 |
| GO:CC | early phagosome | GO:0032009 | 0.066156435 |
| GO:CC | intrinsic component of synaptic membrane | GO:0099240 | 0.067585178 |

Table S4. Gene ontology analysis for genes associated with differentially methylated cytosines overlapping promoter regions or gene bodies.

| **Sex-associated DMCs** (across all or within 34.5˚C individuals) | | | |
|---|---|---|---|
| Source | GO term name | GO term ID | Benjamini-Hochberg FDR |
| GO:MF | phosphatidylinositol phospholipase C activity | GO:0004435 | 0.046729221 |
| GO:MF | phospholipase C activity | GO:0004629 | 0.046729221 |
| GO:MF | ionotropic glutamate receptor activity | GO:0004970 | 0.091253906 |
| GO:MF | glutamate receptor activity | GO:0008066 | 0.091253906 |
| GO:MF | signaling receptor activity | GO:0038023 | 0.091253906 |
| GO:MF | molecular transducer activity | GO:0060089 | 0.091253906 |
| GO:BP | system development | GO:0048731 | 4.67E-05 |
| GO:BP | multicellular organism development | GO:0007275 | 9.97E-05 |
| GO:BP | anatomical structure development | GO:0048856 | 9.97E-05 |
| GO:BP | developmental process | GO:0032502 | 9.97E-05 |
| GO:BP | nervous system development | GO:0007399 | 9.97E-05 |
| GO:BP | animal organ development | GO:0048513 | 0.000461474 |
| GO:BP | cellular developmental process | GO:0048869 | 0.00103919 |
| GO:BP | modulation of excitatory postsynaptic potential | GO:0098815 | 0.001098492 |
| GO:BP | cell differentiation | GO:0030154 | 0.001444742 |
| GO:BP | multicellular organismal process | GO:0032501 | 0.001444742 |
| GO:BP | cell development | GO:0048468 | 0.002087083 |
| GO:BP | glutamate receptor signaling pathway | GO:0007215 | 0.002290445 |
| GO:BP | generation of neurons | GO:0048699 | 0.002290445 |
| GO:BP | neurogenesis | GO:0022008 | 0.002356907 |
| GO:BP | cell junction organization | GO:0034330 | 0.003451143 |
| GO:BP | synapse assembly | GO:0007416 | 0.003451143 |
| GO:BP | establishment of cell polarity | GO:0030010 | 0.003600573 |
| GO:BP | synapse organization | GO:0050808 | 0.004147013 |
| GO:BP | anatomical structure morphogenesis | GO:0009653 | 0.004455126 |
| GO:BP | cell adhesion | GO:0007155 | 0.006689459 |
| GO:BP | biological adhesion | GO:0022610 | 0.006689459 |
| GO:BP | neuron differentiation | GO:0030182 | 0.006689459 |
| GO:BP | cell junction assembly | GO:0034329 | 0.006689459 |
| GO:BP | movement of cell or subcellular component | GO:0006928 | 0.006689459 |
| GO:BP | regulation of postsynaptic membrane potential | GO:0060078 | 0.00887211 |
| GO:BP | positive regulation of nervous system development | GO:0051962 | 0.009089329 |
| GO:BP | regulation of endothelial cell migration | GO:0010594 | 0.009089329 |
| GO:BP | endothelial cell migration | GO:0043542 | 0.011361955 |
| GO:BP | modulation of chemical synaptic transmission | GO:0050804 | 0.019360829 |
| GO:BP | plasma membrane bounded cell projection morphogenesis | GO:0120039 | 0.019360829 |
| GO:BP | regulation of trans-synaptic signaling | GO:0099177 | 0.019360829 |
| GO:BP | neuron projection morphogenesis | GO:0048812 | 0.019360829 |
| GO:BP | circulatory system development | GO:0072359 | 0.019360829 |
| GO:BP | cell projection morphogenesis | GO:0048858 | 0.019360829 |

| Source | GO term name | GO term ID | Benjamini-Hochberg FDR |
|---|---|---|---|
| GO:BP | regulation of membrane potential | GO:0042391 | 0.019360829 |
| GO:BP | system process | GO:0003008 | 0.019554585 |
| GO:BP | neuron development | GO:0048666 | 0.021614364 |
| GO:BP | locomotion | GO:0040011 | 0.021697904 |
| GO:BP | ionotropic glutamate receptor signaling pathway | GO:0035235 | 0.02197245 |
| GO:BP | axon development | GO:0061564 | 0.02197245 |
| GO:BP | ligand-gated ion channel signaling pathway | GO:1990806 | 0.02197245 |
| GO:BP | positive regulation of excitatory postsynaptic potential | GO:2000463 | 0.02197245 |
| GO:BP | cell part morphogenesis | GO:0032990 | 0.023978992 |
| GO:BP | nervous system process | GO:0050877 | 0.023978992 |
| GO:BP | blood vessel morphogenesis | GO:0048514 | 0.023978992 |
| GO:BP | regulation of nervous system development | GO:0051960 | 0.023978992 |
| GO:BP | regulation of cell junction assembly | GO:1901888 | 0.025475418 |
| GO:BP | neuron recognition | GO:0008038 | 0.027444083 |
| GO:BP | regulation of plasma membrane bounded cell projection organization | GO:0120035 | 0.027444083 |
| GO:BP | regulation of nervous system process | GO:0031644 | 0.027444083 |
| GO:BP | protein sulfation | GO:0006477 | 0.027444083 |
| GO:BP | regulation of synapse organization | GO:0050807 | 0.030946139 |
| GO:BP | cellular component morphogenesis | GO:0032989 | 0.035379731 |
| GO:BP | cell morphogenesis involved in neuron differentiation | GO:0048667 | 0.035379731 |
| GO:BP | axonogenesis | GO:0007409 | 0.037290698 |
| GO:BP | neuron projection organization | GO:0106027 | 0.037290698 |
| GO:BP | regulation of synapse assembly | GO:0051963 | 0.037665841 |
| GO:BP | branching involved in mammary gland duct morphogenesis | GO:0060444 | 0.03869607 |
| GO:BP | regulation of cell projection organization | GO:0031344 | 0.040092606 |
| GO:BP | regulation of synapse structure or activity | GO:0050803 | 0.040931059 |
| GO:BP | cell projection organization | GO:0030030 | 0.048480681 |
| GO:BP | cell recognition | GO:0008037 | 0.052433926 |
| GO:BP | excitatory postsynaptic potential | GO:0060079 | 0.052479576 |
| GO:BP | establishment or maintenance of cell polarity | GO:0007163 | 0.056766179 |
| GO:BP | regulation of neuron projection development | GO:0010975 | 0.056766179 |
| GO:BP | neuron projection development | GO:0031175 | 0.056766179 |
| GO:BP | plasma membrane bounded cell projection organization | GO:0120036 | 0.057437684 |
| GO:BP | positive regulation of neurogenesis | GO:0050769 | 0.057437684 |
| GO:BP | regulation of regulated secretory pathway | GO:1903305 | 0.063430303 |
| GO:BP | chemical synaptic transmission, postsynaptic | GO:0099565 | 0.064017036 |
| GO:BP | neuron projection guidance | GO:0097485 | 0.064017036 |
| GO:BP | axon guidance | GO:0007411 | 0.064017036 |
| GO:BP | tube morphogenesis | GO:0035239 | 0.064017036 |
| GO:BP | neuromuscular process controlling balance | GO:0050885 | 0.067510989 |
| GO:BP | vocalization behavior | GO:0071625 | 0.06884488 |
| GO:BP | cell motility | GO:0048870 | 0.06884488 |

| Source | GO term name | GO term ID | Benjamini-Hochberg FDR |
|---|---|---|---|
| GO:BP | localization of cell | GO:0051674 | 0.06884488 |
| GO:BP | regulation of exocytosis | GO:0017157 | 0.06884488 |
| GO:BP | cell surface receptor signaling pathway | GO:0007166 | 0.069936321 |
| GO:BP | blood vessel development | GO:0001568 | 0.072488886 |
| GO:BP | tube development | GO:0035295 | 0.072488886 |
| GO:BP | vasculature development | GO:0001944 | 0.072707903 |
| GO:BP | central nervous system development | GO:0007417 | 0.072707903 |
| GO:BP | cell migration | GO:0016477 | 0.073692838 |
| GO:BP | mammary gland branching involved in pregnancy | GO:0060745 | 0.073942135 |
| GO:BP | negative regulation of toll-like receptor signaling pathway | GO:0034122 | 0.075190437 |
| GO:BP | regulation of system process | GO:0044057 | 0.076765406 |
| GO:BP | postsynapse organization | GO:0099173 | 0.076765406 |
| GO:BP | positive regulation of endothelial cell migration | GO:0010595 | 0.076765406 |
| GO:BP | positive regulation of blood vessel endothelial cell migration | GO:0043536 | 0.076765406 |
| GO:BP | regulation of epithelial cell migration | GO:0010632 | 0.085419205 |
| GO:BP | brain development | GO:0007420 | 0.085419205 |
| GO:BP | action potential | GO:0001508 | 0.088182096 |
| GO:BP | neuromuscular process | GO:0050905 | 0.090126529 |
| GO:BP | tissue migration | GO:0090130 | 0.092197216 |
| GO:BP | syncytium formation by plasma membrane fusion | GO:0000768 | 0.09630765 |
| GO:BP | positive regulation of cell projection organization | GO:0031346 | 0.09630765 |
| GO:BP | cell-cell fusion | GO:0140253 | 0.09630765 |
| GO:BP | chemotaxis | GO:0006935 | 0.096934946 |
| GO:BP | cell morphogenesis involved in differentiation | GO:0000904 | 0.096934946 |
| GO:BP | regulation of cellular component biogenesis | GO:0044087 | 0.096934946 |
| GO:BP | neuron cell-cell adhesion | GO:0007158 | 0.09719147 |
| GO:BP | postsynapse assembly | GO:0099068 | 0.09719147 |
| GO:CC | anchored component of membrane | GO:0031225 | 5.28E-05 |
| GO:CC | cell periphery | GO:0071944 | 0.000783107 |
| GO:CC | neuron projection | GO:0043005 | 0.001236169 |
| GO:CC | cell projection | GO:0042995 | 0.001236169 |
| GO:CC | plasma membrane | GO:0005886 | 0.001236169 |
| GO:CC | neuronal cell body | GO:0043025 | 0.001240491 |
| GO:CC | plasma membrane bounded cell projection | GO:0120025 | 0.001564383 |
| GO:CC | receptor complex | GO:0043235 | 0.001807446 |
| GO:CC | cell junction | GO:0030054 | 0.002505658 |
| GO:CC | cell body | GO:0044297 | 0.002505658 |
| GO:CC | somatodendritic compartment | GO:0036477 | 0.003609509 |
| GO:CC | dendrite | GO:0030425 | 0.005212251 |
| GO:CC | dendritic tree | GO:0097447 | 0.005251196 |
| GO:CC | cell-cell junction | GO:0005911 | 0.008041091 |
| GO:CC | axon | GO:0030424 | 0.017198764 |
| GO:CC | synapse | GO:0045202 | 0.019393832 |
| GO:CC | intrinsic component of membrane | GO:0031224 | 0.021794879 |

| Source | GO term name | GO term ID | Benjamini-Hochberg FDR |
|---|---|---|---|
| GO:CC | external encapsulating structure | GO:0030312 | 0.022018343 |
| GO:CC | extracellular matrix | GO:0031012 | 0.022018343 |
| GO:CC | intrinsic component of plasma membrane | GO:0031226 | 0.022018343 |
| GO:CC | postsynaptic membrane | GO:0045211 | 0.023984963 |
| GO:CC | neuron to neuron synapse | GO:0098984 | 0.025952643 |
| GO:CC | asymmetric synapse | GO:0032279 | 0.025952643 |
| GO:CC | integral component of plasma membrane | GO:0005887 | 0.035700497 |
| GO:CC | cell surface | GO:0009986 | 0.035700497 |
| GO:CC | postsynaptic density | GO:0014069 | 0.035700497 |
| GO:CC | muscle cell projection | GO:0036194 | 0.035700497 |
| GO:CC | muscle cell projection membrane | GO:0036195 | 0.035700497 |
| GO:CC | CNTFR-CLCF1 complex | GO:0097059 | 0.035700497 |
| GO:CC | synaptic membrane | GO:0097060 | 0.035700497 |
| GO:CC | anchoring junction | GO:0070161 | 0.040184607 |
| GO:CC | cytoskeleton | GO:0005856 | 0.045271109 |
| GO:CC | potassium channel complex | GO:0034705 | 0.047773629 |
| GO:CC | sodium channel complex | GO:0034706 | 0.047773629 |
| GO:CC | cation channel complex | GO:0034703 | 0.060547769 |
| GO:CC | postsynaptic specialization | GO:0099572 | 0.063844236 |
| GO:CC | plasma membrane region | GO:0098590 | 0.063844236 |
| GO:CC | postsynaptic specialization membrane | GO:0099634 | 0.063844236 |
| GO:CC | kainate selective glutamate receptor complex | GO:0032983 | 0.078195412 |
| GO:CC | cytolytic granule | GO:0044194 | 0.078195412 |
| GO:CC | cytoplasmic region | GO:0099568 | 0.079007217 |
| GO:CC | postsynapse | GO:0098794 | 0.080518304 |
| GO:CC | plasma membrane protein complex | GO:0098797 | 0.088151107 |
| GO:CC | microfibril | GO:0001527 | 0.094177952 |
| **Universal sex-associated DMCs** (across all individuals) | | | |
| Source | GO term name | GO term ID | Benjamini-Hochberg FDR |
| GO:CC | CNTFR-CLCF1 complex | GO:0097059 | 0.005657706 |
| **Temperature-associated DMCs** | | | |
| Source | GO term name | GO term ID | Benjamini-Hochberg FDR |
| GO:BP | system development | GO:0048731 | 0.034970995 |
| GO:BP | signaling | GO:0023052 | 0.090335271 |
| GO:BP | movement of cell or subcellular component | GO:0006928 | 0.093572399 |
| GO:BP | regulation of signaling | GO:0023051 | 0.098392685 |
| GO:BP | cell migration | GO:0016477 | 0.098392685 |
| GO:BP | regulation of cell communication | GO:0010646 | 0.098392685 |
| GO:BP | multicellular organism development | GO:0007275 | 0.098392685 |
| GO:BP | signal transduction | GO:0007165 | 0.098392685 |
| GO:BP | cell communication | GO:0007154 | 0.098392685 |
| GO:BP | vasculature development | GO:0001944 | 0.098392685 |
| GO:BP | blood vessel development | GO:0001568 | 0.098392685 |
| GO:BP | cellular response to stimulus | GO:0051716 | 0.098392685 |

| Source | GO term name | GO term ID | Benjamini-Hochberg FDR |
|---|---|---|---|
| GO:CC | cell junction | GO:0030054 | 0.008213443 |
| GO:CC | cell periphery | GO:0071944 | 0.008213443 |
| GO:CC | plasma membrane | GO:0005886 | 0.032772209 |

Table S5. Model coefficients for cytosines selected to be included in the predictive model of sex.

| seqnames | starts | ends | coefficients |
|---|---|---|---|
| NW_017707593.1 | 91913 | 91913 | -0.0356773 |
| NW_017707836.1 | 1654499 | 1654499 | 0.03212573 |
| NW_017707977.1 | 543656 | 543656 | 0 |
| NW_017708230.1 | 3394 | 3394 | -0.0089074 |
| NW_017708561.1 | 39079 | 39079 | 0.01909979 |
| NW_017708944.1 | 151 | 151 | -0.011266 |
| NW_017709083.1 | 3729449 | 3729449 | -0.0122084 |
| NW_017709336.1 | 14554887 | 14554887 | -0.0086496 |
| NW_017709336.1 | 2808872 | 2808872 | 0.04981291 |
| NW_017709809.1 | 4507591 | 4507591 | -0.0263089 |
| NW_017709848.1 | 131158 | 131158 | 0 |
| NW_017710078.1 | 1083668 | 1083668 | -0.0258123 |
| NW_017710078.1 | 1172445 | 1172445 | 0 |
| NW_017710621.1 | 24066900 | 24066900 | 0.02225572 |
| NW_017710865.1 | 13502 | 13502 | 0.03607998 |
| NW_017710965.1 | 10634 | 10634 | 0.06808733 |
| NW_017712138.1 | 20423104 | 20423104 | -0.0342119 |
| NW_017712461.1 | 405501 | 405501 | 0.01471531 |
| NW_017713645.1 | 39474677 | 39474677 | -0.0278214 |
| NW_017713709.1 | 15783590 | 15783590 | -0.023263 |
| NW_017713972.1 | 4410508 | 4410508 | 0.0220656 |
| NW_017714183.1 | 13535814 | 13535814 | -0.0080856 |
| NW_017714267.1 | 48405077 | 48405077 | -0.0087841 |
| **intercept** | | | -3.3731516 |

Table S6. Model coefficients for the cytosines selected to be included in the predictive model of incubation temperature.

| seqnames | starts | ends | coefficients |
|---|---|---|---|
| NW_017707537.1 | 1156821 | 1156821 | 0 |
| NW_017707575.1 | 664576 | 664576 | 0 |
| NW_017707593.1 | 31566915 | 31566915 | 0 |
| NW_017707680.1 | 97612 | 97612 | 0 |
| NW_017707689.1 | 612408 | 612408 | 0 |
| NW_017707789.1 | 246112 | 246112 | 0 |
| NW_017707836.1 | 13872742 | 13872742 | 0 |
| NW_017707888.1 | 59383 | 59383 | 0 |
| NW_017707888.1 | 6175388 | 6175388 | 0 |
| NW_017707901.1 | 19373924 | 19373924 | 0 |
| NW_017707901.1 | 20157398 | 20157398 | 0 |
| NW_017707947.1 | 55173633 | 55173633 | 0 |
| NW_017707977.1 | 2267140 | 2267140 | -0.0124876 |
| NW_017708049.1 | 7444667 | 7444667 | 0 |
| NW_017708054.1 | 4751706 | 4751706 | 0 |
| NW_017708115.1 | 890584 | 890584 | 0 |
| NW_017708254.1 | 362681 | 362681 | -0.0163736 |
| NW_017708254.1 | 522064 | 522064 | 0 |
| NW_017708373.1 | 2095 | 2095 | 0 |
| NW_017708483.1 | 12588 | 12588 | 0 |
| NW_017708551.1 | 15179 | 15179 | 0 |
| NW_017708624.1 | 12784596 | 12784596 | 0 |
| NW_017708624.1 | 15345883 | 15345883 | 0 |
| NW_017708672.1 | 2204425 | 2204425 | 0 |
| NW_017708718.1 | 50920 | 50920 | -0.0036008 |
| NW_017708734.1 | 52212 | 52212 | 0 |
| NW_017708765.1 | 4254832 | 4254832 | 0 |
| NW_017708765.1 | 5603885 | 5603885 | 0 |
| NW_017708776.1 | 109935 | 109935 | 0 |
| NW_017708796.1 | 556167 | 556167 | 0 |
| NW_017708899.1 | 10669901 | 10669901 | 0 |
| NW_017708899.1 | 14520000 | 14520000 | 0 |
| NW_017708899.1 | 15794844 | 15794844 | 0 |
| NW_017709017.1 | 1002275 | 1002275 | 0 |
| NW_017709017.1 | 729544 | 729544 | 0 |
| NW_017709029.1 | 144476 | 144476 | 0 |
| NW_017709042.1 | 22962869 | 22962869 | 0 |
| NW_017709065.1 | 706150 | 706150 | 0 |
| NW_017709065.1 | 800711 | 800711 | 0 |
| NW_017709067.1 | 31571 | 31571 | 0 |
| NW_017709154.1 | 15587 | 15587 | 0 |
| NW_017709210.1 | 101550 | 101550 | 0 |
| NW_017709210.1 | 151409 | 151409 | 0 |
| NW_017709336.1 | 13952461 | 13952461 | 0 |

| seqnames | starts | ends | coefficients |
|---|---|---|---|
| NW_017709344.1 | 528603 | 528603 | 0 |
| NW_017709480.1 | 8165125 | 8165125 | 0 |
| NW_017709599.1 | 10555075 | 10555075 | 0 |
| NW_017709599.1 | 11694301 | 11694301 | 0 |
| NW_017709599.1 | 11803781 | 11803781 | 0 |
| NW_017709599.1 | 7424511 | 7424511 | -0.0014596 |
| NW_017709599.1 | 8690707 | 8690707 | 0 |
| NW_017709854.1 | 27383056 | 27383056 | 0 |
| NW_017709984.1 | 695115 | 695115 | 0.00080832 |
| NW_017710057.1 | 260299 | 260299 | 0 |
| NW_017710078.1 | 1172445 | 1172445 | 0 |
| NW_017710164.1 | 90031 | 90031 | 0 |
| NW_017710321.1 | 10692071 | 10692071 | 0 |
| NW_017710432.1 | 3336829 | 3336829 | 0 |
| NW_017710448.1 | 397324 | 397324 | 0 |
| NW_017710448.1 | 821723 | 821723 | 0 |
| NW_017710466.1 | 1741 | 1741 | -5.62E-06 |
| NW_017710504.1 | 52087 | 52087 | 0 |
| NW_017710534.1 | 5333531 | 5333531 | -0.003189 |
| NW_017710560.1 | 331268 | 331268 | -0.0006225 |
| NW_017710621.1 | 23656719 | 23656719 | 0 |
| NW_017710746.1 | 4471645 | 4471645 | 0 |
| NW_017710946.1 | 3320418 | 3320418 | 0 |
| NW_017710965.1 | 10634 | 10634 | 0.02114739 |
| NW_017710976.1 | 756238 | 756238 | 0.04154243 |
| NW_017711041.1 | 3475323 | 3475323 | 0 |
| NW_017711054.1 | 18358 | 18358 | 0 |
| NW_017711096.1 | 25640420 | 25640420 | 6.32E-05 |
| NW_017711096.1 | 28717706 | 28717706 | 0 |
| NW_017711096.1 | 5619169 | 5619169 | 0 |
| NW_017711201.1 | 646956 | 646956 | 0 |
| NW_017711293.1 | 1811343 | 1811343 | 0 |
| NW_017711293.1 | 464681 | 464681 | 0 |
| NW_017711319.1 | 5863921 | 5863921 | 0 |
| NW_017711344.1 | 7021 | 7021 | 0 |
| NW_017711346.1 | 244563 | 244563 | 0 |
| NW_017711487.1 | 10068264 | 10068264 | -0.0051853 |
| NW_017711487.1 | 2051686 | 2051686 | 0 |
| NW_017711487.1 | 4940281 | 4940281 | 0 |
| NW_017711487.1 | 5148758 | 5148758 | 0 |
| NW_017711527.1 | 11969347 | 11969347 | 0 |
| NW_017711567.1 | 1335170 | 1335170 | 0 |
| NW_017711568.1 | 10926618 | 10926618 | 0 |
| NW_017711568.1 | 18109023 | 18109023 | -0.0020448 |
| NW_017711692.1 | 15736498 | 15736498 | 0 |
| NW_017711861.1 | 5666 | 5666 | 0 |
| NW_017711921.1 | 1469 | 1469 | 0 |

| seqnames | starts | ends | coefficients |
|---|---|---|---|
| NW_017711921.1 | 1474 | 1474 | 0 |
| NW_017711985.1 | 207462 | 207462 | 0 |
| NW_017712118.1 | 16197695 | 16197695 | 0 |
| NW_017712138.1 | 116740 | 116740 | 0 |
| NW_017712138.1 | 20423104 | 20423104 | 0 |
| NW_017712138.1 | 8576097 | 8576097 | 0.01340404 |
| NW_017712138.1 | 959869 | 959869 | 0 |
| NW_017712186.1 | 6018016 | 6018016 | -0.0139219 |
| NW_017712194.1 | 160546 | 160546 | 0 |
| NW_017712303.1 | 2366038 | 2366038 | 0 |
| NW_017712464.1 | 9557375 | 9557375 | 0 |
| NW_017712924.1 | 3348061 | 3348061 | 0 |
| NW_017713026.1 | 1944 | 1944 | 0 |
| NW_017713083.1 | 108585 | 108585 | 0 |
| NW_017713083.1 | 1856586 | 1856586 | 0 |
| NW_017713165.1 | 343 | 343 | 0 |
| NW_017713334.1 | 62391 | 62391 | 0 |
| NW_017713446.1 | 6101896 | 6101896 | 0 |
| NW_017713475.1 | 1947470 | 1947470 | 0 |
| NW_017713485.1 | 320211 | 320211 | 0 |
| NW_017713513.1 | 11479708 | 11479708 | 0.01415717 |
| NW_017713513.1 | 2852287 | 2852287 | 0 |
| NW_017713516.1 | 444921 | 444921 | 0 |
| NW_017713629.1 | 17031206 | 17031206 | 0.01315481 |
| NW_017713629.1 | 18243123 | 18243123 | 0 |
| NW_017713629.1 | 18712057 | 18712057 | 0.00178697 |
| NW_017713629.1 | 19278690 | 19278690 | 0 |
| NW_017713629.1 | 19583010 | 19583010 | 0.01283963 |
| NW_017713645.1 | 17492529 | 17492529 | 0 |
| NW_017713645.1 | 9641978 | 9641978 | 0 |
| NW_017713697.1 | 15320021 | 15320021 | 0.0047776 |
| NW_017713697.1 | 20082957 | 20082957 | 0 |
| NW_017713697.1 | 29698341 | 29698341 | 0 |
| NW_017713697.1 | 4576647 | 4576647 | 0 |
| NW_017713700.1 | 1069506 | 1069506 | 0.00399025 |
| NW_017713703.1 | 317477 | 317477 | 0 |
| NW_017713709.1 | 1234746 | 1234746 | 0 |
| NW_017713709.1 | 2137652 | 2137652 | 0 |
| NW_017713709.1 | 21601852 | 21601852 | 0 |
| NW_017713709.1 | 9212745 | 9212745 | 0 |
| NW_017713772.1 | 18699692 | 18699692 | 0 |
| NW_017713772.1 | 6328963 | 6328963 | 0.00382647 |
| NW_017713851.1 | 22425584 | 22425584 | 0 |
| NW_017713894.1 | 2814789 | 2814789 | 0 |
| NW_017713955.1 | 3014 | 3014 | 0 |
| NW_017713998.1 | 7077131 | 7077131 | 0 |
| NW_017714045.1 | 91353 | 91353 | 0 |

| seqnames | starts | ends | coefficients |
|---|---|---|---|
| NW_017714045.1 | 9230558 | 9230558 | 0.00635438 |
| NW_017714054.1 | 463859 | 463859 | 0 |
| NW_017714106.1 | 1217490 | 1217490 | 0 |
| NW_017714180.1 | 1648 | 1648 | 0 |
| NW_017714194.1 | 16670684 | 16670684 | 0 |
| NW_017714219.1 | 89786 | 89786 | 0 |
| NW_017714267.1 | 42786365 | 42786365 | 0 |
| NW_017714267.1 | 49033156 | 49033156 | 0 |
| NW_017714270.1 | 352270 | 352270 | 0 |
| NW_017714436.1 | 1312872 | 1312872 | 0 |
| NW_017714440.1 | 542 | 542 | 0 |
| NW_017714442.1 | 193621 | 193621 | 0.0122612 |
| NW_017714443.1 | 762448 | 762448 | 0 |
| NW_017714467.1 | 842 | 842 | 0 |
| **intercept** | | | 2.40E+01 |

Table S7. Sequencing summary for reduced-representation sequencing libraries from embryonic gonads.

| Sample | Temperature | Clutch | Total reads | Total aligned reads | Percent alignment | Bisulfite conversion efficiency |
|---|---|---|---|---|---|---|
| 1 | 33.5 | WO-03 | 5929216 | 3395703 | 57.2706914 | 99.9297 |
| 3 | 34.5 | WO-10 | 11151190 | 6374060 | 57.1603569 | 99.5941 |
| 4 | 32 | WO-01 | 14163866 | 7761664 | 54.7990499 | 99.5042 |
| 5 | 29 | WO-14 | 14515134 | 7901948 | 54.4393734 | 99.1275 |
| 6 | 29 | WO-03 | 8532667 | 4656977 | 54.578211 | 99.9642 |
| 7 | 33.5 | WO-01 | 11466703 | 6132469 | 53.4806648 | 99.6724 |
| 8 | 33.5 | WO-14 | 10666274 | 5542201 | 51.9600472 | 99.3716 |
| 9 | 29 | WO-14 | 8563375 | 4797032 | 56.0180069 | 99.9326 |
| 10 | 33.5 | WO-14 | 5772967 | 3079819 | 53.3489798 | 99.2703 |
| 11 | 32 | WO-14 | 8346295 | 4493569 | 53.8390867 | 99.5219 |
| 12 | 33.5 | WO-10 | 8423048 | 4524574 | 53.7165881 | 99.4297 |
| 13 | 34.5 | WO-14 | 6072181 | 3444702 | 56.7292378 | 99.9121 |
| 14 | 34.5 | WO-08 | 6577901 | 3781251 | 57.4841579 | 98.6111 |
| 15 | 32 | WO-14 | 7360671 | 3942372 | 53.559954 | 99.8255 |
| 16 | 33.5 | WO-08 | 6166782 | 3310883 | 53.6889905 | 99.8789 |
| 17 | 29 | WO-08 | 8729572 | 4972378 | 56.960158 | 99.8904 |
| 18 | 29 | WO-10 | 7859923 | 4401316 | 55.9969353 | 99.2152 |
| 19 | 34.5 | WO-01 | 8430493 | 4806315 | 57.0110787 | 99.7643 |
| 21 | 32 | WO-08 | 8992885 | 4957061 | 55.1220326 | 99.5529 |
| 22 | 32 | WO-03 | 8967757 | 4990198 | 55.6459993 | 99.8441 |
| 23 | 29 | WO-01 | 7878311 | 4475609 | 56.809245 | 99.8505 |
| 24 | 32 | WO-10 | 8438401 | 4815033 | 57.0609645 | 99.6923 |

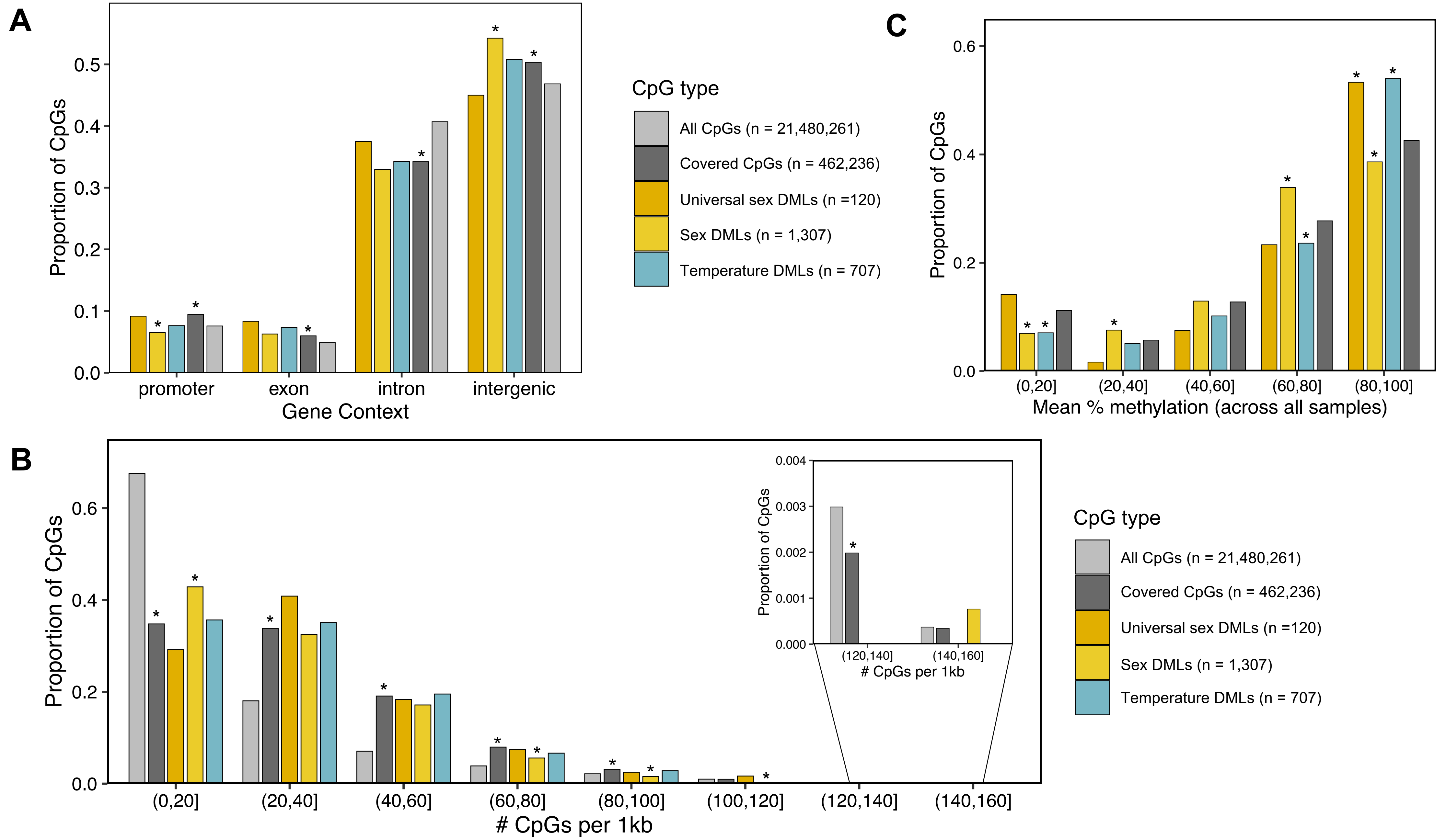


**Figure S1.** Genomic characterization of sex- and temperature-associated differentially methylated loci. (A) Proportion of CpGs overlapping promoters, exons, introns, and intergenic regions. (B) Proportion of CpGs overlapping 1kb tiles of varying CpG density. (C) Proportion of CpGs with varying mean percent methylation across all samples. Asterisks indicate significant difference in the proportion of DMCs in a context category compared to the covered CpGs or significant difference in the proportion of covered CpGs in a context category compared to all the CpGs in the alligator genome at an FDR threshold of 10%.

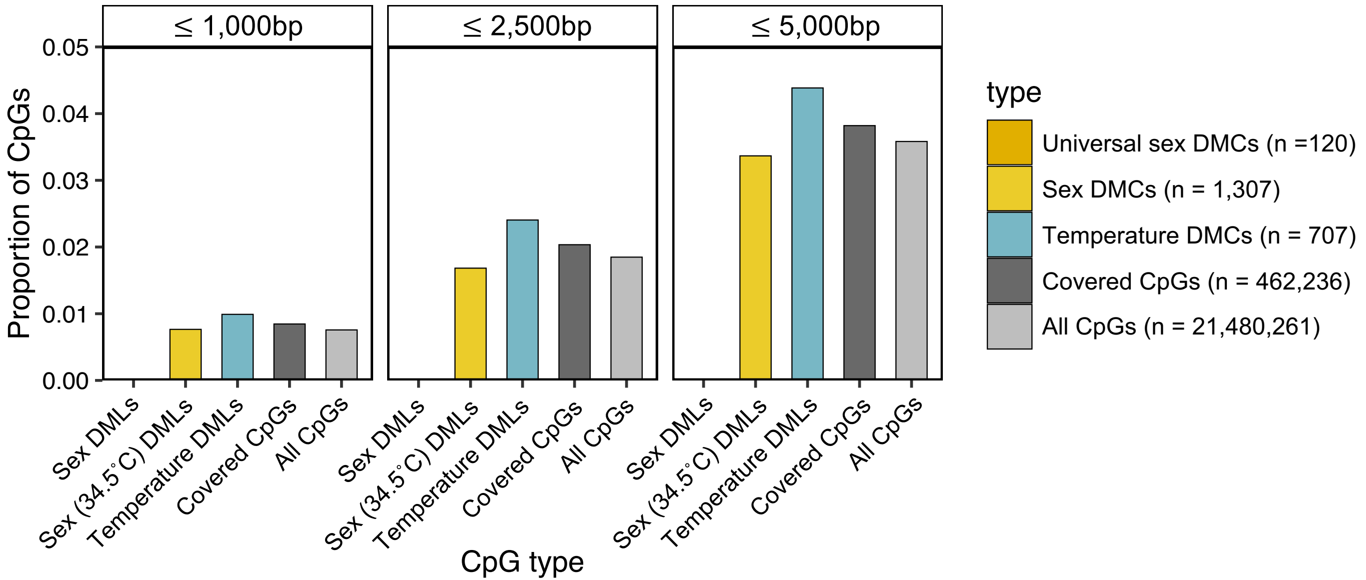


**Figure S2.** Associations between differentially methylated loci and putative estrogen-response elements. Proportion of CpGs within 1kb, 2.5kb, and 5kb of putative estrogen-response element. Asterisks indicate significant difference in the proportion of covered CpGs in a context category compared to all the CpGs in the alligator genome at an FDR threshold of 10%.

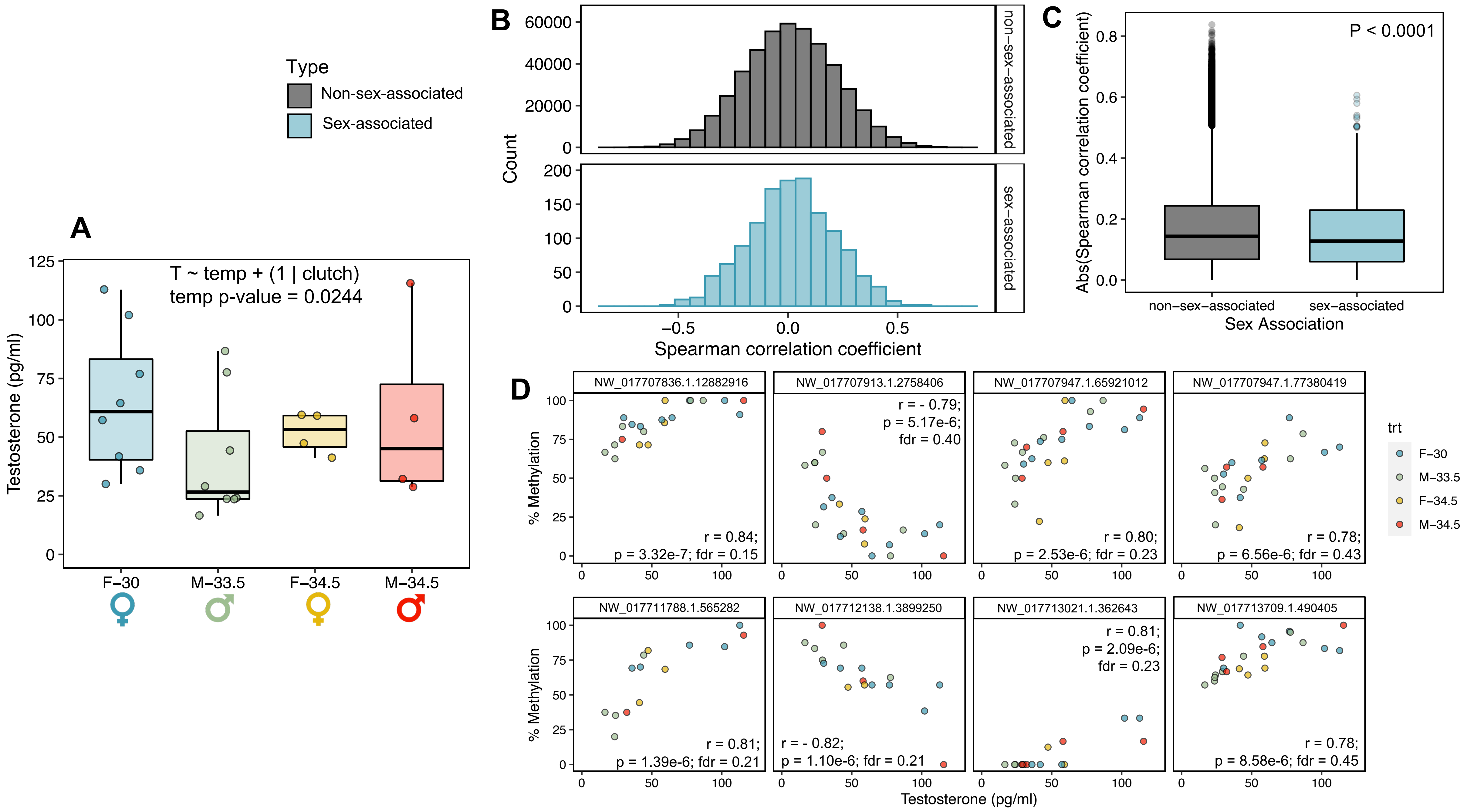


**Figure S3.** Correlations between plasma testosterone (T) concentrations and genome-wide DNA methylation patterns. (A) Boxplot comparing plasma T concentrations between treatment groups. (B) Histograms of Spearman correlation coefficients for T correlations, with panels separating loci not identified as sex-associated via pairwise comparisons (top) from loci identified as sex-associated differentially methylated cytosines (DMCs; bottom). (C) Boxplot comparing absolute Spearman correlation coefficients between sex-associated DMCs and all other loci. Sex-associated loci collectively exhibited a significantly lower absolute Spearman correlation with plasma T compared to loci not identified as sex-associated. (C) Scatterplots depicting the relationship between percent methylation and plasma T concentration for the loci exhibiting the top 8 most significant correlations. For boxplots, central line depicts median, box depicts interquartile range (IQR), vertical lines depict the maximum and minimum values, and values greater or less than 1.5*IQR are depicted as points.

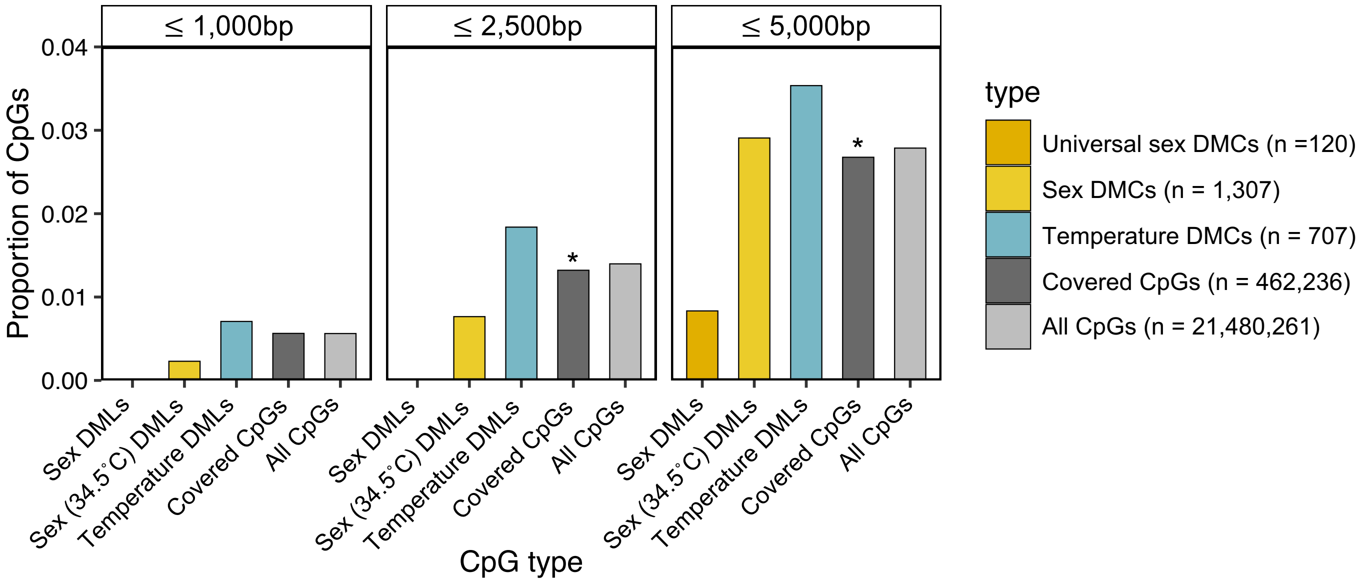


**Figure S4.** Associations between differentially methylated loci and putative androgen-response elements. Proportion of CpGs within 1kb, 2.5kb, and 5kb of putative androgen-response element. Asterisks indicate significant difference in the proportion of covered CpGs in a context category compared to all the CpGs in the alligator genome at an FDR threshold of 10%.
